# Specific detection and deletion of the Sigma-1 receptor in neurons and glial cells for functional characterization *in vivo*

**DOI:** 10.1101/2022.06.08.494880

**Authors:** Qing Liu, Qilin Guo, Li-Pao Fang, Honghong Yao, Anja Scheller, Frank Kirchhoff, Wenhui Huang

## Abstract

The chaperon protein sigma-1 receptor (S1R) has been discovered over forty years ago. Recent pharmacological studies using S1R exogenous ligands demonstrated a promising therapeutical potential of targeting the S1R for several neurological disorders. Although intensive *in vitro* studies have revealed S1Rs are mainly residing at the membrane of the endoplasmic reticulum (ER), the cell-specific *in vivo* expression pattern of S1Rs is still unclear, mainly due to the lack of a reliable detection method which also prevented a comprehensive functional analysis.

Here, first, we identified a highly specific antibody using S1R knockout (KO) mice and established an immunohistochemical protocol involving a 1% SDS antigen retrieval step. Second, we characterized the S1R expression in the mouse brain and can demonstrate that the S1R is widely expressed: in principal neurons, interneurons, and all glial cell types. Finally, we generated a novel Cre-dependent S1R conditional KO mouse (S1R flox) to study cell type-specific functions of the S1R. As a proof of concept, we successfully ablated S1R expressions in neurons or microglia employing neuronal and microglial Cre-expressing mice, respectively. In summary, we provide powerful tools to cell-specifically detect, delete and functionally characterize S1R *in vivo*.

## Introduction

The sigma-1 receptor (S1R) is a chaperon protein primarily residing at mitochondria-associated membranes (MAM) of the endoplasmic reticulum (ER) with a single membrane-spanning domain, which is considered as a pluripotent modulator involved in many aspects of cellular functions (Su, Su, Nakamura, & Tsai, 2016). Previous studies by immunohistochemistry and mRNA expression profiling experiments suggested that S1Rs are highly expressed in the central nervous system (CNS) and can also be detected in other organs such as liver, kidney, and muscles (Couly et al., 2022; Su et al., 2016; Zhang et al., 2014). However, it is still hard to conclude the cellular sources of S1R in the CNS due to seemingly paradoxical results from immunohistochemistry. For instance, using a custom-made antibody Alonso et al. detected S1Rs only in neurons in the brain and spinal cord, while Palacios et al., using an independently custom-made antibody could observe S1R expression also in OLs (Alonso et al., 2000; Palacios et al., 2003), and Ruscher et al., using a commercial antibody observed S1R immunoreactivity co-localized with the cytoskeleton indicated by glial fibrillary acidic protein (GFAP) as well as with the galactocerebroside-enriched membrane microdomains of reactive astrocytes in the peri-infarct area of rat brains after cerebral stroke (Ruscher et al., 2011). Of note, it is not clear whether the specificity of the antibodies used in the aforementioned studies was tested by S1R KO mice/cell lines as rigorous controls. To date, only one custom-made antibody against S1R (termed Ab^Ruoho^) generated by Arnold Ruoho’s group was validated by S1R KO mice, showing high specificity for immunohistochemistry in the brain and spinal cord (though it did not work well for immunoblot) (Mavlyutov et al., 2016; Mavlyutov, Epstein, Andersen, Ziskind-Conhaim, & Ruoho, 2010; Nakamura et al., 2019). However, in these studies the fine structures of Ab^Ruoho^ stained cells in brain slices were not displayed with high magnification, neither were co-immunostainings combined for different cell type markers with Ab^Ruoho^ performed. Therefore, it is hard to verify the detailed expression pattern of S1Rs in neurons and glial cells *in vivo*.

Neurons and glial cells interact with each other to orchestrate diverse CNS functions. Previous studies using constitutive S1R KO mice suggest S1Rs are involved in the maintenance of cognitive, psychiatric, and motor functions, particularly with aging (Couly et al., 2022). Moreover, the S1R is considered as an enigmatic therapeutic target for various neurological disorders upon activation by its exogenous ligands including agonists and antagonists (Sałaciak & Pytka, 2022; Schmidt & Kruse, 2019). However, the contribution of cell-type-specific S1Rs to modulate the neural network activity under physiological and pathological conditions as well as upon activation is still not well understood, largely due to the lack of research tools inducing S1R deletion/overexpression in targeted cells *in vivo*.

In the current work, we prepared tissue lysates of brains and spinal cords from WT and S1R KO mice to screen six commercial antibodies against the S1R for their immuno-specificity by immunoblot. We obtained one rabbit monoclonal antibody (#61994, Cell Signaling) displaying very high specificity for S1R by Western blot analysis. We further revealed that after the antigen retrieval using 1% sodium dodecyl sulfate (SDS), this antibody demonstrated highly specific immunolabelling of S1Rs *in situ* in the CNS. Combining immunostaining for different cell type markers, we identified that S1Rs were widely expressed in the CNS, i.e. in principal neurons, interneurons, astrocytes, oligodendrocyte precursor cells (OPCs), OLs, and microglia. In addition, we found that unlike previous reports (Francardo et al., 2014; Ruscher et al., 2011), S1R expression in astrocytes was not correlated with the GFAP-labelled cytoskeleton, neither in healthy brain slices nor after acute brain injuries. Secondly, we generated a S1R flox mouse with exons 1-3 of *Sigmar1* (gene name of S1R, also called *Oprs1*) flanked by two loxP sites. By cross-breeding this S1R flox mouse with two Cre-driver mouse lines targeting neurons and microglia respectively, we were able to show the specific deletion of S1Rs in targeted cells *in vivo*. Taken together, we introduce a reliable protocol to detect S1Rs by immunoblotting as well as by immunohistochemistry. We also provide a novel S1R flox mouse for cell type-specific ablation of S1Rs *in vivo*. In the future, these tools will facilitate the functional analysis of S1Rs *in vivo*.

## Materials and methods

### Animals

All mice used in this study were maintained at the animal facility of the CIPMM in a temperature-(22°C ± 2°C) and humidity-controlled facility with a 12-h light/dark cycle. Animal husbandry and procedures were performed at the animal facility of CIPMM, University of Saarland according to European and German guidelines for the welfare of experimental animals. Animal experiments were approved by the Saarland state’s ‘‘Landesamt für Gesundheit und Verbraucherschutz” in Saarbrücken/Germany (animal license number:34/2016, 36/2016, 03/2021 and 08/2021).

For antibody testing, 4 to 8 weeks old mice of either sex were used in this study. *Sigmar1* global knockout (S1R KO) mice were generated by GemPharmatech (Nanjing, China) by deleting the entire encoding region (∼10359bp) of *Sigmar1*.

RiboTag mice (Rlp22^HA^) (Sanz et al., 2009) were introduced to immunoprecipitate ribosome-associated translated mRNA in targeted glial cells upon breeding with different glia-specific Cre-driver mice. Specifically, GLAST-CreERT2 mice for astrocytes (Mori et al., 2006), CX3CR1-CreERT2 mice for microglia (Yona et al., 2013), and NG2-CreERT2 mice for OPCs (W. Huang et al., 2014) were used.

The Sigmar1 flox (S1R^fl/fl^) mouse line was generated through the “Dalmatian Mouse Action” of GemPharmatech (Nanjing, China). CRISPR/Cas9 technology was used to modify the *Sigmar1* gene (*Oprs1*). Briefly, single guide RNA (sgRNA) was transcribed *in vitro*, and the donor vector containing exons 1-3 of *Sigmar1* flanked by two loxP sites was constructed. Cas9, sgRNA and the donor vector were microinjected into the fertilized eggs of C57BL/6J mice. sgRNA directed Cas9 endonuclease cleavage at about 6 kb upstream of exon1 and downstream of 3’UTR and create a double strand break (DSB). The following primer sequences were used for genotyping PCR (forward: 5’-AAG CAG AAG AGC AGC TAG TGC TG-3’, reverse: 5’-TGA GAC AGG GTT TCT CTG TAT AGC C-3’).

To obtain cell type-specific S1R knockout mice, S1R^fl/fl^ mice were crossed to NEX-Cre mice (Goebbels et al., 2006) to induce the specific knockout of S1Rs in principal neurons within the dorsal telencephalon and hippocampus or crossed to CX3CR1-CreERT2 mice (Yona et al., 2013) to induce the knockout of S1Rs in microglia upon tamoxifen administration.

### Tamoxifen Induction

Tamoxifen (CC99648, Carbobution) was dissolved in Miglyol (3274, Caesar & Loretz, Hilden) at a concentration of 10 mg/ml. All mice crossed with CreERT2-driver mouse lines were injected intraperitoneally with tamoxifen (100 mg/kg of body weight) for five consecutive days at the age of 4 weeks.

### Stab wound injury (SWI) model

Adult mice (10 weeks old) were used for stab wound injuries (SWI) as described before with some modifications (W. Huang, Bai, Meyer, & Scheller, 2020). Briefly, under isoflurane anesthesia, animals were fixed in a stereotaxic frame with a heat plate. After sterile cleaning and skin incision, a 2 mm cranial grove was drilled in the right neocortex at Bregma from 0.5-2.5 mm, lateral 1.5 mm. A sterile razor blade (2 mm width) was inserted vertically into brain parenchyma (1 mm deep) parallel with the middle line. The lesion was cleaned and closed with sutures. Animals were postsurgically injected subcutaneously with analgesic and antiphlogistic agents for three consecutive days. After 3 days post injury (3 dpi), mice were deeply anesthetized and perfused with 4% paraformaldehyde (PFA) dissolved in 0.1 M phosphate buffer (PB, PH 7.4). Coronal sections (35 µm) were collected and used for immunostaining.

### Magnetic-associated cell sorting (MACS) of glial cells

MACS was performed according to the manufacturer’s instruction (Miltenyi Biotec) with some modifications as shown previously (Fang et al., 2022). In brief, 4 weeks old mice were perfused with cold Hank’s balanced salt solution without Ca^2+^ and Mg^2+^ (HBSS, H6648, Gibco) and cortices were dissected in ice cold HBSS. After the removal of debris (130-107-677, Miltenyi Biotec), cells were resuspended with 1 mL “re-expression medium” containing NeuroBrew-21 (1:50 in MACS neuro Medium) (130-093-566 and 130-093-570, Miltenyi Biotec) and 200 mM L-glutamine (1:100, G7513, Sigma) at 37°C for 30 min.

For OPC sorting, cells were incubated with Fc-receptor blocker (provided with the CD140 microbeads kit) for 10 min at 4°C, followed by a 15 min incubation with 10 µL microbeads mixture containing antibodies directed against CD140 (130-101-502, Miltenyi Biotec), NG2 (130-097-170, Miltenyi Biotec) and O4 (130-096-670, Miltenyi Biotec) in 1:1:1 at 4°C.

For sorting of astrocytes, microbeads containing antibodies directed against ACSA-2 (130-097-678, Miltenyi Biotec) were used.

For microglia sorting, microbeads containing antibodies directed against CD11b (130-093-634, Miltenyi Biotec) were used.

### Western blot analysis

After anesthesia with 1 mg/kg ketamine and 0.5 mg/kg xylazine, mice were transcardially perfused with ice-cold PBS. The dorsal region of the cortex was dissected from coronal brain slices (1 mm). Segments from the cervical spinal cord segment were collected. Specimen were stored at -80°C until tested. RIPA lysis buffer (89900, Thermo Scientific) containing 1x protease inhibitor cocktail (05892970001, Roche) was used to extract protein. Protein concentration was measured using the Bicinchoninic Acid (BCA) assay kit (Thermo Fisher Scientific). After adding 1x protein loading buffer (42526.01, SERVA) containing 5% ß-Mercaptoethanol (M6250, Sigma-Aldrich), protein samples were denatured 5 min at 95°C. Equal amounts of lysates (10-30 µg) of each mouse were separated by 10% SDS-polyacrylamide gel electrophoresis (PAGE, 43289.01, SERVA) and transferred onto nitrocellulose (NC) membranes (QP0907015, neoLab). Homogeneous protein-transfer onto NC membranes was evaluated by Ponceau S staining. After blocking with 5% non-fat milk powder (A0830,0500, PanReac AppliChem,) in 1x PBS for 1 h at room temperature (RT), NC membranes were incubated with primary antibodies at 4°C overnight (**Table 1**) in TBST solution (Tris-base buffer with 0.1% Tween-20). The next day, membranes were washed three times with TBST and incubated with corresponding horseradish peroxidase (HRP) conjugated secondary antibodies (**Table 2**) in TBST for 1 h at RT. For detecting different proteins on the same NC membrane, the previous antibodies were stripped off by stripping buffer for 20 min and then incubated with other primary antibodies.

**Table 1.** Primary antibodies used for Western blot and immunohistochemistry.

| Antigen | Host Species | Antibody type | Source | Catalog# | Dilution |
| --- | --- | --- | --- | --- | --- |
| Sigma-1 receptor | Rabbit | monoclonal | Cell Signaling | #61994 | WB: 1:1000, IHC: 1:500 |
| Sigma-1 receptor | Rabbit | monoclonal | Cell Signaling | #74807 | WB: 1:1000, IHC: 1:500 |
| Sigma-1 receptor | Mouse | monoclonal | Santa Cruz | sc-137075 | WB: 1:500, IHC: 1:500 |
| Sigma-1 receptor | Rabbit | polyclonal | Invitrogen | #42-3300 | WB: 1:500, IHC: 1:500 |
| Sigma-1 receptor | Rabbit | polyclonal | Abcam | Ab53852 | WB: 1:500, IHC: 1:500 |
| Sigma-1 receptor | Rabbit | polyclonal | Proteintech | 15168-1-AP | WB: 1:500, IHC: 1:500 |
| CD11b | Rabbit | monoclonal | Abcam | Ab133357 | WB: 1:1000 |
| GFAP | Rabbit | Polyclonal | Dako | Z 0334 | WB: 1:1000 |
| GAPDH | Mouse | monoclonal | Sigma-Aldrich | G8795 | WB: 1:1000 |
| α-Tubulin | Mouse | monoclonal | Sigma-Aldrich | T6074 | WB: 1:1000 |
| Glutamate Synthase (GS) | Mouse | monoclonal | BD | 610518 | IHC: 1:500 |
| GFAP | Goat | Polyclonal | Abcam | Ab53554 | IHC: 1:500 |
| PDGFRα | Goat | Polyclonal | R&D Systems | AF1042 | IHC: 1:500 |
| Quaking 7 (for APC CC-1) | Mouse | monoclonal | Calbiochem | OP80 | IHC: 1:200 |
| Iba1 | Goat | Polyclonal | Abcam | ab5076 | IHC: 1:500 |
| S100 beta (S100B) | Mouse | Monoclonal | Abcam | ab66028 | IHC: 1:500 |
| Calbindin-D28K | Mouse | Monoclonal | Sigma-Aldrich | C9848 | IHC: 1:500 |
| NeuN | Mouse | monoclonal | Millipore | MAB377 | IHC: 1:500 |
| Myelin Basic Protein (MBP) | Mouse | monoclonal | Biolegend | SMI99 | IHC: 1:500 |
| Parvalbumin | Mouse | monoclonal | Sigma-Aldrich | P3088 | IHC: 1:500 |
| Somatostatin | Rat | monoclonal | Millipore | MAB354 | IHC: 1:500 |
| α-Actinin | Mouse | monoclonal | Sigma-Aldrich | A7811 | IHC: 1:500 |

**Table 2.** Secondary antibodies used for Western blot and immunohistochemistry.

| Antibody | Source | Dilution |
| --- | --- | --- |
| goat anti-rabbit IgG HRP | Dianova, 111-035-045 | WB: 1: 5000 |
| goat anti-mouse IgG HRP | Sigma, A9044 | WB: 1: 10000 |
| Alexa Fluor 488 donkey anti-mouse | Invitrogen | IHC: 1: 1000 |
| Alexa Fluor 488 donkey anti-goat | Thermo Fisher, A11055 | IHC: 1: 1000 |
| Alexa Fluor 546 donkey anti-mouse | Thermo Fisher, A10036 | IHC: 1: 1000 |
| Alexa Fluor 647 donkey anti-rabbit | Thermo Fisher, A31573 | IHC: 1: 1000 |
| DAPI (stain for nuclei) | Biochimica, A10010010 | IHC: 1: 1000 |

For MACS-purified cells, 40 µl RIPA lysis buffer (89900, Thermo Scientific) and the equal amount of 1x loading buffer with 5% ß-Mercaptoethanol were added per sample. After denaturation for 5 min at 95°C, 10 µl of each protein sample were separated by SDS-PAGE and assessed by Western blot as described above. All primary antibodies are listed in **Table 1** and secondary antibodies in **Table 2**, respectively.

The immunoblots were processed using enhanced chemiluminescence (ECL) reagent (541015, Biozym) and ChemiDoc Imaging System (BioRad). The immunoblot intensity was quantified with ImageJ software (ImageJ 1.53f51, NIH. USA).

### Immunohistochemistry

After anesthesia, mice were transcardially perfused with 5 ml PBS and followed with 15 ml 4% PFA. Dissected brains were post-fixed in 4% PFA at 4°C overnight. Then, coronal brain sections or transverse spinal sections were prepared by vibratome (VT1000S, Leica). The regular free-floating immunostaining was performed as previously described (W. Huang et al., 2020). Two new protocols with antigen retrieval (AR) were established for S1R staining with the antibody #61994:

AR^SDS^ protocol for immunostaining of S1Rs in the brain: brain slices were pre-treated with 1% SDS (CN30.1, Roth) in 1x PBS for 10 min at RT for antigen retrieval (Brown et al., 1996). After three times washing with 1x PBS, the blocking buffer (1x Fish Gelatin Blocking Agent (22010, BioTium), 0.5% Triton x-100 in PBS) was added to slices and incubated for 40 min at RT to decrease background signal. All primary antibodies were diluted in blocking buffer.

AR^EtOH-SDS^ protocol for S1R immunostaining in the spinal cord: spinal slices were incubated with 100% ethanol (EtOH) at 4°C overnight with gentle shake for delipidation. After washing with 1x PBS, 1% SDS pre-treatment was performed as described in AR^SDS^ protocol above.

Brain and spinal cord slices were incubated at 4°C for 2 nights with primary antibodies at appropriate dilutions as shown in **Table 1**. After washing with 1x PBS, sections were incubated with corresponding secondary antibodies (**Table 2**) for 2 hours at RT. DAPI was used for nuclear staining. After washing three times with 1x PBS, slices were mounted with Immu-Mount (9990402, Thermo) (Figure 2A).

### Ribosome immunoprecipitation (IP)

After perfusion with ice-cold HBSS, cortical samples were dissected from mouse brain and stored at −80°C until use. Tissues were homogenized in ice-cold lysis buffer (50 mM Tris, pH 7.4, 100 mM KCl, 12 mM MgCl_2_, 1% NP-40, 1 mM DTT, 1x protease inhibitor, 200 units/ml RNasin (Promega) and 0.1 mg/ml cycloheximide (Sigma-Aldrich) in RNase-free deionized H_2_O) 10% w/v with homogenizer (Precellys 24, PeQlab). Homogenates were centrifuged at 10,000 g at 4°C for 10 min to remove cell debris. Supernatants were collected, from which 50 μl were removed for input analysis. Anti-HA Ab (1:100, # MMS-101P, Covance) was added to the supernatant and slowly rotated at 4°C. Protein G-Dynabeads (Thermo Fisher Scientific) were equilibrated with lysis buffer by washing three times. After 4 h of incubation with HA Ab, 100 μl pre-equilibrated beads were added to each sample and incubated overnight at 4°C. After 10-12 h, samples were washed with high-salt buffer (50 mM Tris, 300 mM KCl, 12 mM MgCl_2_, 1% NP-40, 1 mM DTT, 1x protease inhibitor, 100 units/ml RNasin and 0.1 mg/ml cycloheximide in RNase-free deionized H_2_O) three times for 5 min at 4°C. At the end of the washing, beads were magnetized and 150 µl RA1 lysis buffer from NucleoSpin RNA Plus XS Kit (40990.50, Macherey-Nagel) was added to the beads. RNA was extracted followed with manufacturer’s instructions (NucleoSpin RNA Plus XS, Macherey-Nagel).

### Quantitative real time PCR (qPCR)

RNA concentration was determined using NanoDrop from IP and input RNA. 100 µg of RNA was used to synthesize first-strand complementary DNA (cDNA) using Omniscript kit (205113, QIAGEN). qPCR was performed with EvaGreen (27490, Axon) in a CFX96 Real-Time System (BioRad). The standard two-step program was used: 94 °C for 10 min, then 40 cycles at 94 °C for 15 sec, 60 °C for 1 min. The expression of *Sigmar1, Gfap, Itgam*, and *Pdgfra* was measured. For primer see **Table 3**. Relative expression of targeted genes was determined using the ΔΔCt method with normalization to β-actin expression.

**Table 3.**
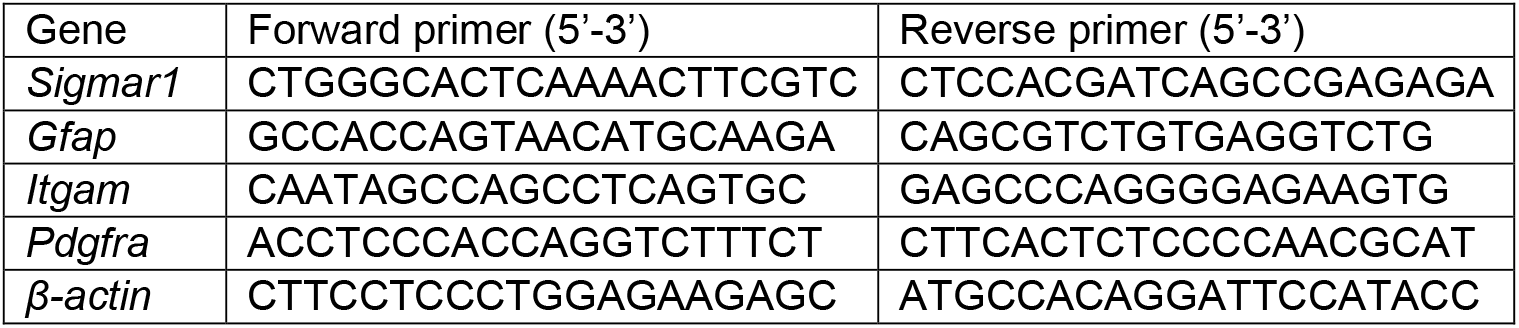
Primers used for qPCR

### Image acquisition and quantification

Images were acquired using an epifluorescence microscope system AxioScan.Z1 (Zeiss, Oberkochen, Germany), with a Plan-Apochromat 20x/0.8 M27 objective and a Zeiss confocal microscope system LSM 880 with Plan-Apochromat 40x/1.3 Oil DIC UV-IR M27 objective (Zeiss, Oberkochen, Germany).

For each immunostaining, two coronal brain sections per mouse were collected randomly at the hippocampus level. Epifluorescence AxionScan images were used for quantification of S1R expression in NeuN^+^ cells using ZEN 3.1 (blue edition) software (Figure 3E). For quantification of S1R expression in interneurons and glial cells (Figure 3F, G; 4E, F; 5E, F; 6E, F; 7E, F) as well as in NeuN^+^ cells in NEX-Cre x S1R^fl/fl^ mice (Figure 10D, E), confocal stacks were taken from six cortical layers and two different areas from corpus callosum. For deletion of S1R in microglia in CX3CR1-CreERT2 x S1R^fl/fl^ mice, confocal images were randomly taken from three areas over the dorsal cortex. Cell counting was performed using ZEN 3.0 SR (black edition) (Carl Zeiss, 16.0.2.306).

Four to five transversal sections per mouse were collected randomly from the cervical spinal cord. Three random areas from the white matter (WM) and grey matter (GM) were taken by confocal microscopy and analyzed by ZEN 3.0 SR (black edition).

### Statistical analysis

Data were analyzed using GraphPad Prism 9.3.1 statistical software (GraphPad, San Diego, CA). All data were given as Mean ± SEM. For statistical analysis, the independent-sample *t*-test was used. *P* values of ≤ 0.05 were considered statistically significant.

## Results

### Specific detection of the S1R by immunoblot

To identify reliable S1R antibodies, we screened six commercially available antibodies on tissue homogenates obtained from the cerebral cortex and spinal cord of WT and S1R KO mice by immunoblotting. The protein samples were prepared by RIPA buffer containing 1% Triton X-100 to release total proteins of the tissue. We loaded 5-30 μg proteins per sample for SDS-PAGE. Prior to incubating with primary antibodies, we stained the blotted nitrocellulose (NC) membranes with Ponceau S solution to confirm proteins of different sizes were uniformly transferred. After incubating the NC membrane with S1R antibodies, we took advantage of the high sensitivity of the HRP (horseradish peroxidase)-based enhanced chemiluminescent (ECL) system to detect S1Rs. To detect potential unspecific signals, we exposed each membrane incubated with different S1R antibodies to a digital imaging system for as long as 15 min. We observed that one monoclonal rabbit antibody #61994 (Ab^#61994^) from Cell Signaling showed strong signals at the expected size of the S1R (25 kD) in WT mice which were completely absent in the KO mice (Figure 1A, right). Even with the long exposure time (15 min), we detected only faint bands at positions of higher molecular weight. In addition, Ab^#61994^ generated the same immunoblot results to detect S1Rs in whole brain lysates (data not shown). Another monoclonal rabbit antibody #74807 from CST did not show any bands at 25 kD but unspecific upper bands in WT and KO mice. The monoclonal mouse antibody sc-137075 from Santa Cruz showed relatively weak but specific bands of S1Rs at 25 kD, in line with previous studies (Moreno et al., 2014; Yang, Shen, Li, Stanford, & Guo, 2020). However, Ab^sc-137075^ also detected many other proteins of different sizes both in WT and KO mice. Other polyclonal rabbit antibodies, i.e. 42-3300 from Invitrogen, ab53852 from Abcam and 15168-1-AP from Proteintech, showed bands at 25 kD and other positions both in WT and KO mice (Ab^15168-1-AP^ showed weaker bands at 25 kD in KO mice), indicating unspecific detections of S1Rs by those antibodies for immunoblot (Figure 1A). Taken together, Ab^#61994^ was identified as the most specific antibody to detect S1Rs in CNS tissues by immunoblot.

**Figure 1.**
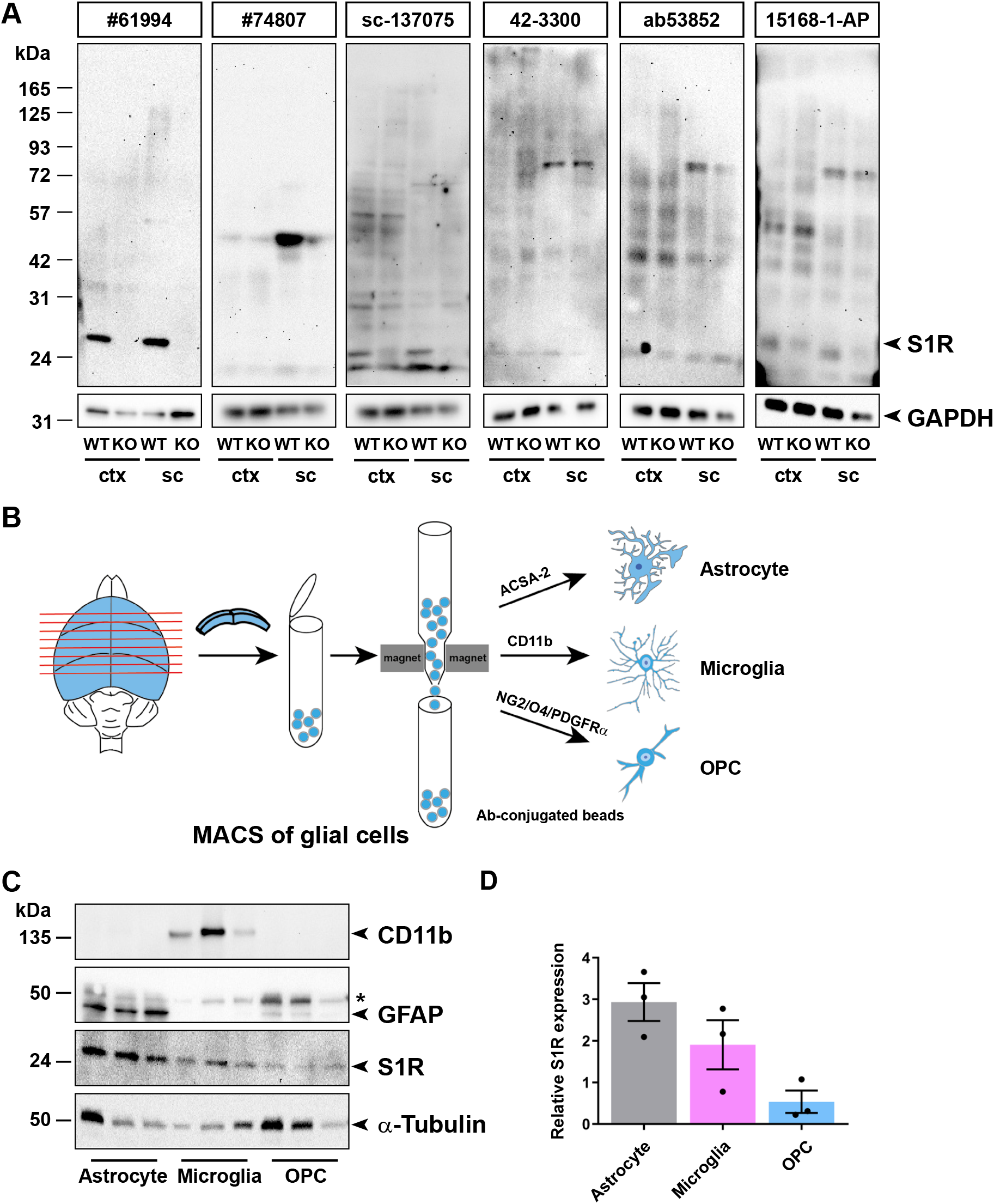
Detection of S1Rs in the CNS by immunoblot. (**A**) Full-length scan of the S1R immunoblot of protein lysates from cortex (ctx) and spinal cord (sc) tissue of WT and S1R KO mice. S1R antibodies (Abs) from Cell Signaling (#61994 and #74807), Santa Cruz (sc-137075), Invitrogen (42-3300), Abcam (ab53852), and Proteintech (15168-1-AP) were used. The correct molecular weight of S1R is 25 kDa. GAPDH was used as loading control. The same membrane was reused for #74807 after stripping off sc-137075. (**B**) Illustration of magnetic-associated cell sorting (MACS) of glial cells from mouse cortex. ACSA-2, CD11b and NG2/O4/PDGFRα conjugated beads were used to purify astrocytes, microglia and OPCs, respectively. (**C**) Immunoblots with expression of S1Rs in sorted astrocytes, microglia and OPCs from mouse brain with CD11b and GFAP immunoblot demonstrating the purity of microglia and astrocyte from MACS, respectively. α-Tubulin was used as loading control. Asterisk (*) indicates the band of α-Tubulin which was not totally stripped off. (**D**) Quantification of grey values of bands from C showing the relative expression of S1R proteins in different glial cells (normalized to α-Tubulin). n = 3 mice.

Previous transcriptome profiling studies using purified cells from postnatal mice demonstrated that *Sigmar1* was widely expressed in neurons and glial cells, and was even detected with higher expression level in microglia and OPCs than other cells (Figure supplement 1A) (Zhang et al., 2014). To better compare *Sigmar1* expression levels in glial cells of adult mice, we purified translated mRNA directly from astrocytes, microglia, and OPCs of adult mouse cerebral cortex using a Cre-dependent RiboTag approach (Figure supplement 1B, C and D). Quantitative PCR (qPCR) results suggested that astrocytes expressed the highest *Sigmar1* translated mRNA level while microglia showed the lowest level (Figure supplement 1E). To further determine the S1R protein expression in adult glia, we purified astrocytes, microglia, and OPCs from the cortex of adult WT mice by magnetic-activated cell sorting (MACS) and performed immunoblots with the specific Ab^#61994^ (Figure 1B). We observed that S1Rs could be detected in protein samples from purified astrocytes, microglia, and OPCs. A comparison using α-tubulin as loading controls demonstrated that the relative S1R protein expression was highest in astrocytes but lowest in OPCs (Figure 1C and D). Thus, glial cells do express S1R proteins with variable levels (astrocytes > microglia > OPCs), however, inconsistent with the translated mRNA levels (astrocytes > OPCs > microglia).

### Establishment of a reliable protocol for S1R immunohistochemistry

To investigate the S1R expression *in situ*, a specific S1R antibody working for immunohistochemistry is highly demanded. Because paraffin or cryo-section preparations are known to be harmful to antigen preservations for immunohistochemistry (Hira et al., 2019; Shi, Cote, & Taylor, 1997), we used free-floating vibratome sections of formaldehyde-fixed brain tissues. We started with the regular protocol working well in our previous studies (W. Huang et al., 2020; Wenhui Huang, Guo, Bai, Scheller, & Kirchhoff, 2019) to test the S1R antibodies (Figure 2A). However, we observed that Ab^#61994^, Ab^#74807^ (data not shown), and Ab^sc-137075^ did not show any labelling in WT and KO mice (Figure 2B and Figure supplement 2A). We also found that Ab^ab53852^ and Ab^15168-1-AP^ showed weak immunostaining in neuron-like cell bodies which wasn identical in WT and KO mice (Figure supplement 2B and C). In addition, Ab^ab53852^ and Ab^15168-1-AP^ strongly labelled many cells with a pattern very similar to anti-GFAP stainings, mostly in corpus callosum and hippocampus of WT and KO mice. We noticed that such GFAP-like staining was the only immuno-labelling of Ab^42-3300^, however, both in WT and KO mice (Figure supplement 2D). Therefore, these antibodies are not suitable for immunohistochemistry and specific immuno-labelling of cells in the mouse brain.

**Figure 2.**
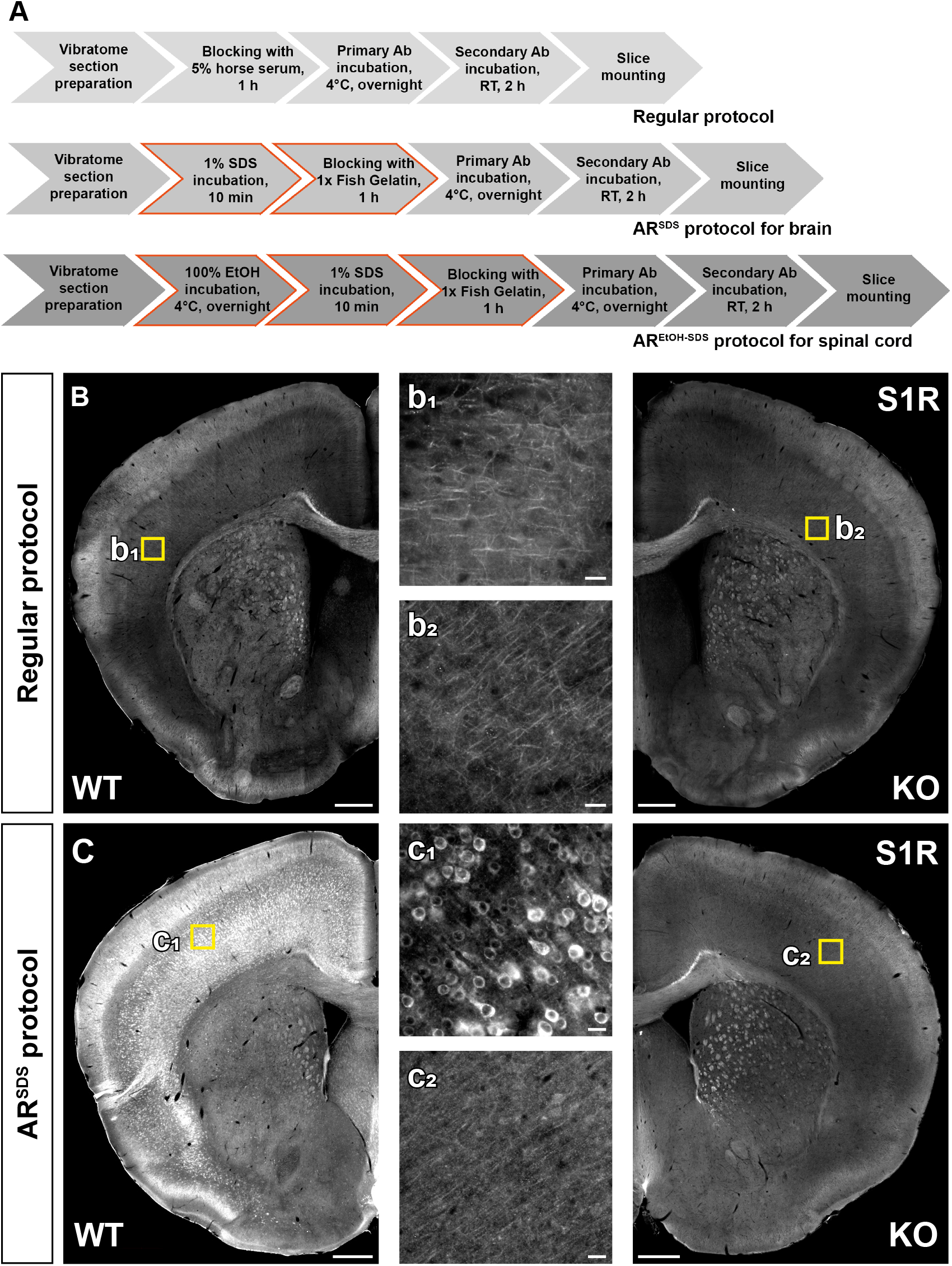
Establishing immunohistochemical protocols to specifically detect S1Rs. **(A)** Immunohistochemical protocols tested for vibratome sections of brain and spinal cord. The regular protocol without antigen retrieval was compared to AR^SDS^ protocol with antigen retrieval (AR) using 1% SDS for brain slices. The modified AR^SDS^ protocol with AR using 100% ethanol and 1% SDS sequentially was used for spinal sections (AR^EtOH-SDS^). (**B-C**) Fluorescent images of S1R immunostainings performed with the regular protocol (B) or AR^SDS^ protocol (C) using Ab^#61994^. Magnified images (b_1_, b_2,_ c_1_, c_2_) showing corresponded yellow boxes in the cortex of WT (b_1_, c_1_) and KO (b_2_, c_2_) mice. Scale bars = 200 µm in A-B, 5 µm in a_1_, a_2_, b_1_, b_2_.

SDS has been suggested as an antigen retrieval (AR) reagent for antibodies detecting denatured proteins in IHC (Brown et al., 1996; Wilson & Bianchi, 1999). Considering that Ab^#61994^ specifically recognized SDS-denatured S1Rs for immunoblot, we thereby treated vibratome brain slices with SDS (1%, 10 min RT) for antigen retrieval prior to the blocking step (Figure 2A). And indeed, we observed bright and clear immunoreactivity to Ab^#61994^ in WT mice which was completely absent in S1R KO mice, strongly indicating the capability of Ab^#61994^ to specifically detect S1Rs in IHC (Figure 2C). Detailed analysis revealed the AR^SDS^ protocol (i.e. 1% SDS for antigen retrieval + Ab^#61994^) specifically detected S1R-expressing cells over the brain, except some unspecifically stained white matter tracts in brain stem and cerebellum (Figure supplement 3B and C). We also noticed that unlike previous reports (Mavlyutov et al., 2010), the cerebral cortex, hippocampus, thalamus, and olfactory bulb area showed strong immuno-labelling of S1Rs by this protocol (Figure supplement 3A, a_1_-a_3_ and B, b_1_-b_3_). Although in images with higher magnifications a background staining of tiny puncta could be seen in WT and KO mice (Figure supplement 3C), Ab^#61994^ immuno-labelling clearly demonstrated the ER-like perinuclear ring structures of the S1R staining as reported in previous studies using EYFP-tagged S1Rs in cultured cells (Hayashi & Su, 2007). In addition, this protocol could immuno-label S1Rs in other organs such as liver (Figure supplement 4A-D) and heart (Figure supplement 4E-H). Therefore, the current AR^SDS^ protocol is able to reliably identify S1R expression *in situ*.

We tested the effect of 1% SDS pre-treatment for other S1R antibodies as well. However, we did not observe improved immunostaining by Ab^#74807^ (data not shown), Ab^sc-137075^, Ab^42-3300^, and Ab^ab53852^ compared to a treatment without SDS (Figure supplement 5A-C). In line with the immunoblot results, the immuno-labelling by Ab^15168-1-AP^ in WT mice was improved after the antigen retrieval, whereas in KO mice similar but weaker immunostaining was also observed (Figure supplement 5D). In addition, the GFAP-like staining could always be found using Ab^42-3300^, Ab^ab53852^, and Ab^15168-1-AP^ in WT and KO mice even after the antigen retrieval (Figure supplement 3B-D). Taken together, except Ab^#61994^, the other commercial S1R antibodies failed to provide a reliable immuno-labelling of S1Rs for IHC.

### S1Rs are expressed in neurons and glial cells in the forebrain

We took advantage of the newly established IHC AR^SDS^ protocol to study the expression of S1Rs in the CNS. We observed similar patterns of S1R immunoreactivity in the brains of mice at different ages from postnatal day 7 (P7) to 24 w (Figure supplement 6A-D). Thus, we performed co-immunostaining for S1Rs and different cell markers in the brain of 8 w old mice to study S1R expression in detail, with particular focus on the cerebral cortex (grey matter) and corpus callosum (cc, white matter) (Fig. 3-7).

We combined NeuN (a pan-neuronal marker) and S1R immunostaining to evaluate the expression of S1Rs in neurons. We found that throughout the forebrain, the majority of neurons were immuno-positive for S1R (Figure 3A and B). For example, in all cortical layers S1R immuno-labelling was found in ∼85% of NeuN^+^ cells in layer 1 (L1) and in more than 90% of NeuN^+^ cells in layer 2-6 (Figure 3E). Since NeuN also labelled interneurons in addition to principal neurons, we further studied the expression of S1Rs in two major interneuron types, i.e. parvalbumin (PV)^+^ and somatostatin (Sst)^+^ interneurons (Figure 3C-D). We found still the majority of PV^+^ or Sst^+^ interneurons were expressing S1R, however, with different proportions. Specifically, the proportion of PV^+^ interneurons immuno-labelled for S1R was ∼70% in L1, ∼80% in L2/3, and ∼90% in L5 and L6 (Figure 3F), while ∼90% of Sst^+^ interneurons showed immunoreactivity of S1R in L2-L6 (Figure 3G). Please note, virtually no PV^+^ or Sst^+^ cells were found in L1.

**Figure 3.**
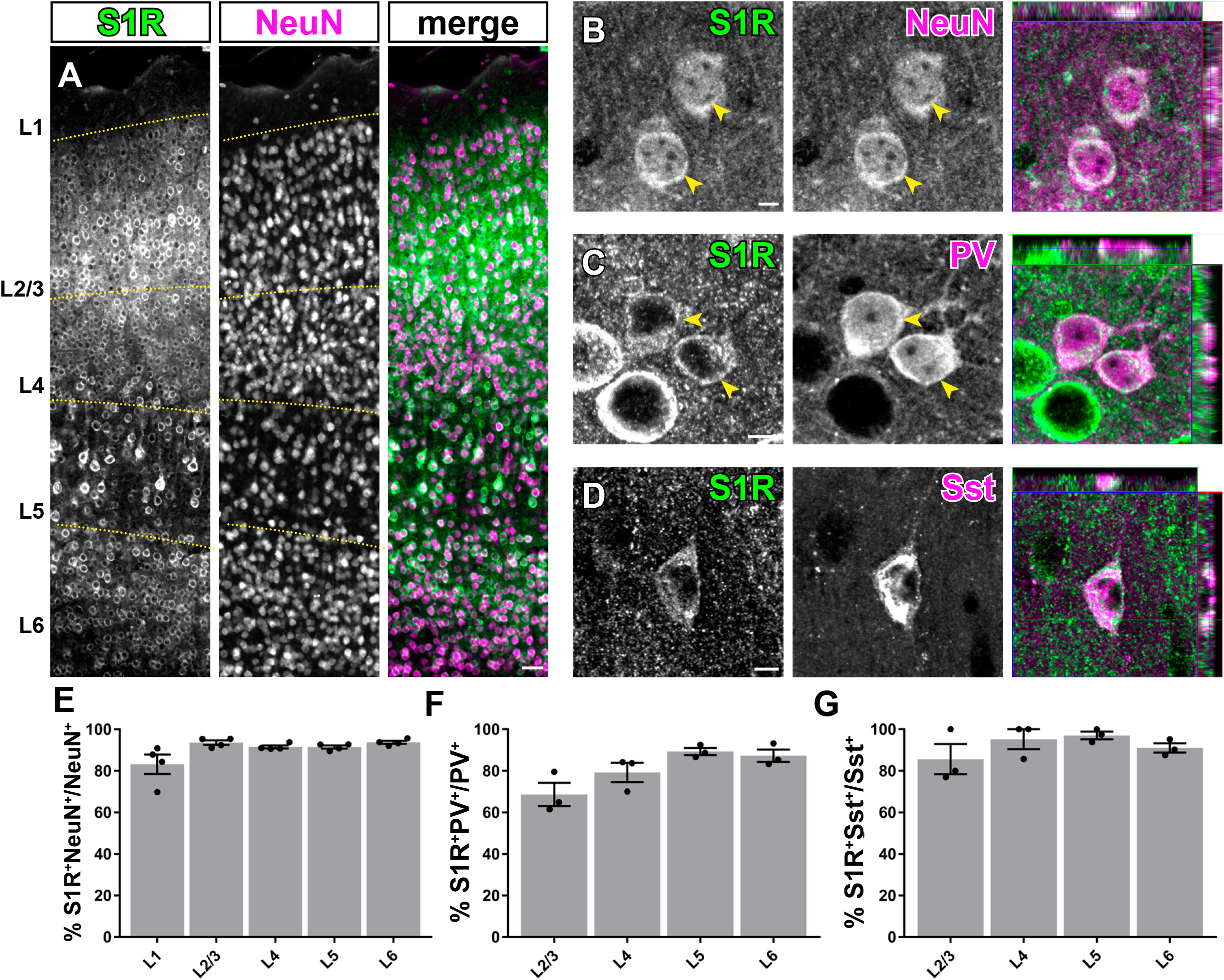
S1Rs are expressed abundantly in neurons. (**A**) Immunohistochemical detection of S1Rs in NeuN^+^ cells in different cortical layers (L1-L6). Almost all NeuN^+^ cells were co-localized with S1R immuno-labelling. (**B**) Confocal images depicting the ring-like structure of S1R immunostaining in NeuN^+^ cells. (**C**-**D**) Detection of S1Rs in Parvalbumin^+^ (PV, C) and Somatostatin^+^ (Sst, D) interneurons. The rightmost images of B-C showing the orthogonal views. (**E**-**G**) The proportions of S1R^+^ cells in NeuN^+^ (E), PV^+^ (F), Sst^+^ neurons from different cortical layers. n = 3 mice. Scale bars = 50 µm in A, 5 µm in B-D.

For detection of astrocytes we used glutamine synthetase (GS) immunostaining. We observed that although the perinuclear ring structure of S1R in GS^+^ cells appeared thinner than in neurons, still most of GS^+^ cells in different areas across the forebrain could be immuno-labelled for S1R (Figure 4A-B). We found ∼80-90% GS^+^ cells expressing S1R in cortical layers and cc (Figure 4E-F). Notably, we did not find an S1R staining in astrocytes that mimicked a typical GFAP-containing cytoskeleton well known from anti-GFAP immunostainings in the healthy brain (Figure 4C-D).

**Figure 4.**
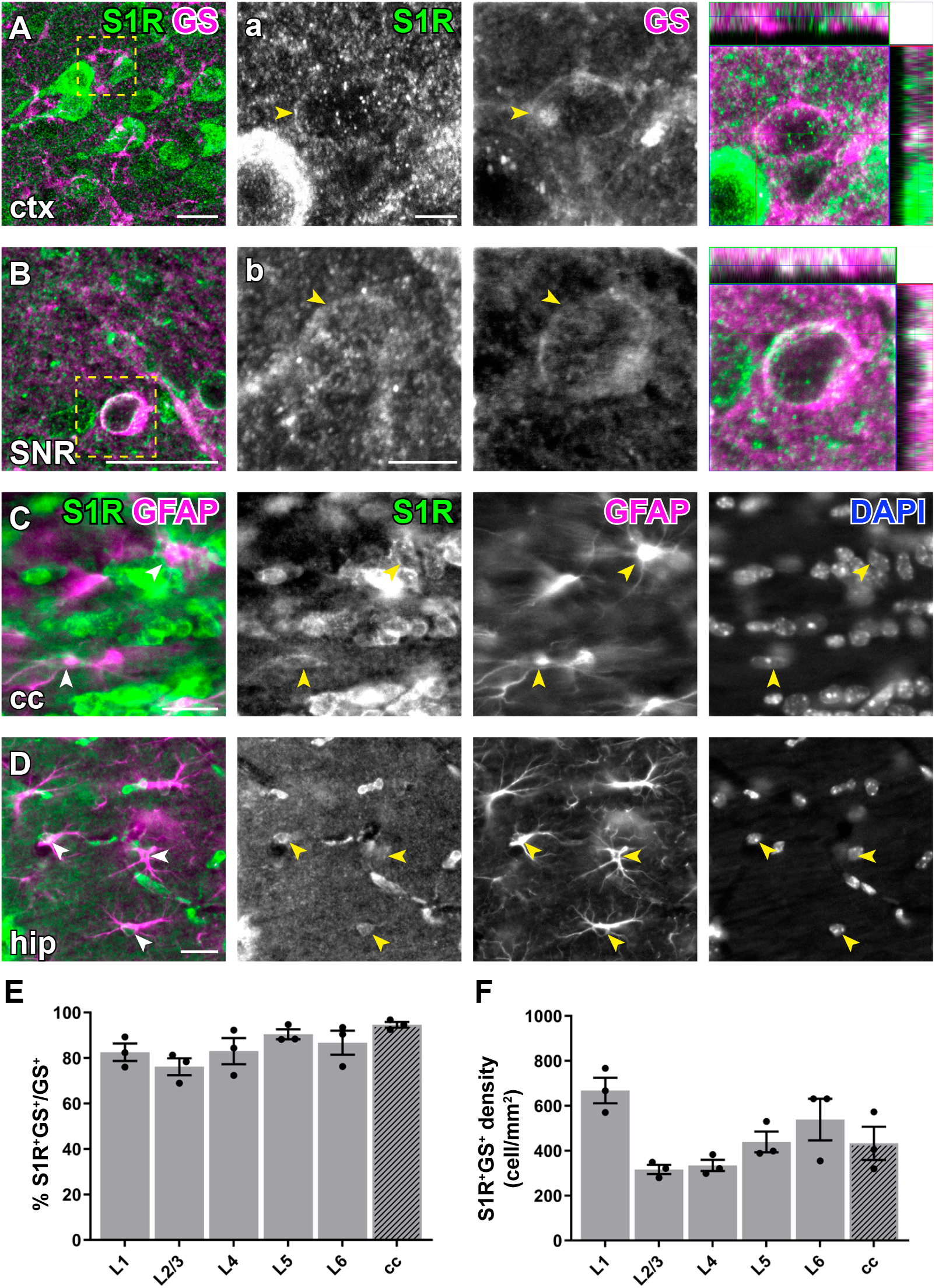
Immunohistochemical detection of S1R expression in forebrain astrocytes. (**A-B**) S1R immunoreactivities in GS^+^ astrocytes in ctx (A), substantia nigra (SNR, B). (**a-c**) Magnified images corresponding to boxed areas in A-B. Arrowhead indicated S1R staining in astrocytes. The orthogonal views are shown in the rightmost pictures. (**C-D**) Axioscan images showing the ring-like structure of S1R staining in GFAP^+^ astrocytes in the corpus callosum (cc, C), hippocampus (hip, D). (**E-F**) Quantification of proportions and densities of GS^+^ cells expressing S1R^+^ in different cortical layers (L1-L6) and cc. n = 3 mice. Scale bars = 20 µm in A-D, 5 µm in a-c.

To study S1R expression in oligodendrocyte lineage cells, we performed PDGFRα (Pα, an OPC marker) and APC CC1 (an OL marker) immunostainings. With the AR^SDS^ protocol, we were able to detect S1R expression in almost all OPCs (Figure 5) and OLs (Figure 6), both in grey matter (e.g. the cortex) and in white matter (e.g. the cc). To our knowledge, this is the first time that S1R expression has been observed in OPCs by immunolabeling.

**Figure 5.**
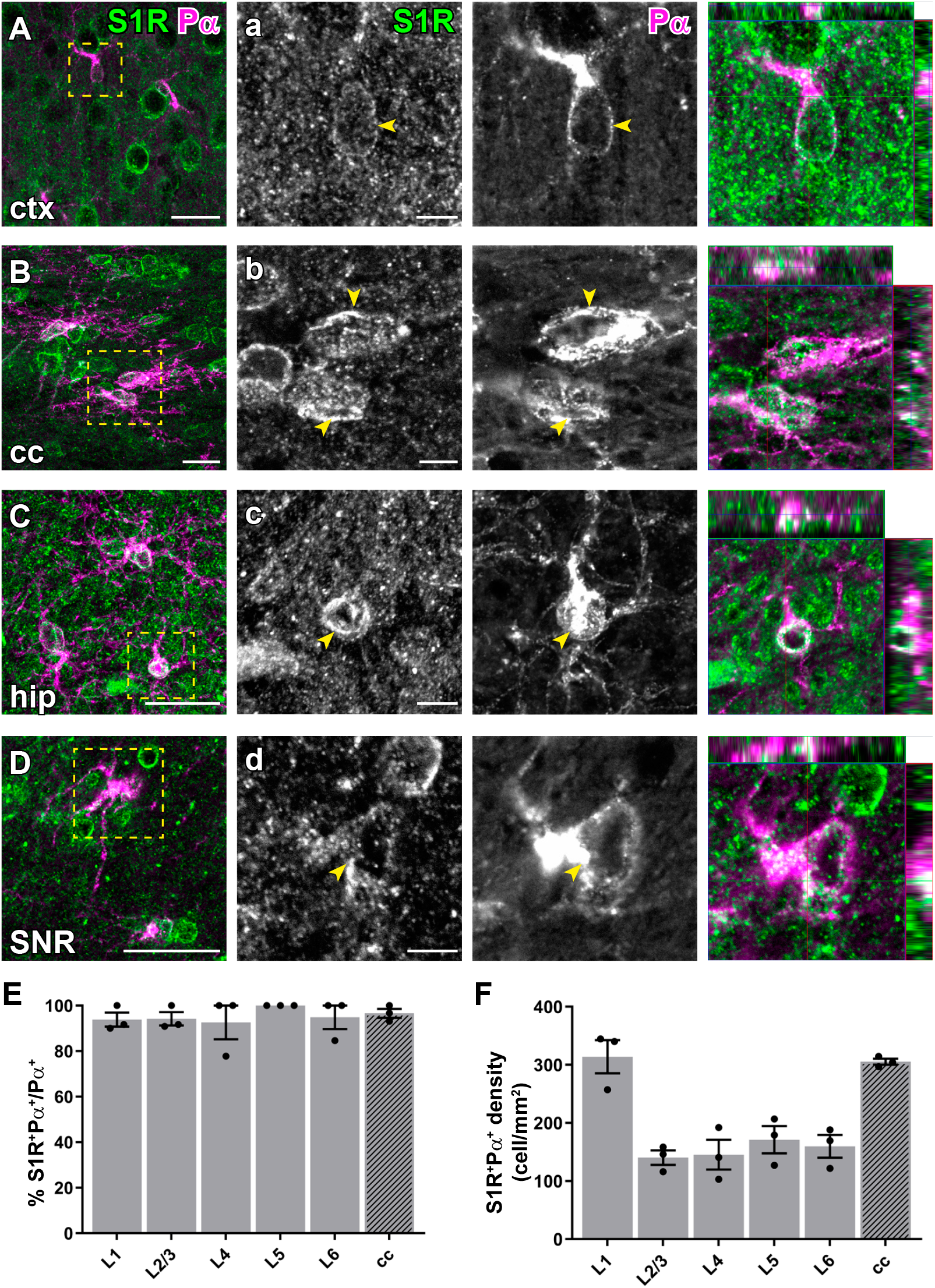
Detection of the S1R expression in OPCs in the forebrain. (**A-D**) S1R immunostaining co-localized with PDGFRα^+^ (Pα) OPCs in ctx (A), cc (B), hip (C), SNR (D). (**a-d**) Magnified images showing boxed areas from A-D. Arrowhead indicated S1R staining in OPCs. The rightmost images indicating the orthogonal views. (**E-F**) Quantification of proportions (E) and densities (F) of S1R^+^Pα^+^cells in different cortical layers (L1-L6) and cc. n = 3 mice. Scale bars = 20 µm in A-D, 5 µm in a-d.

**Figure 6.**
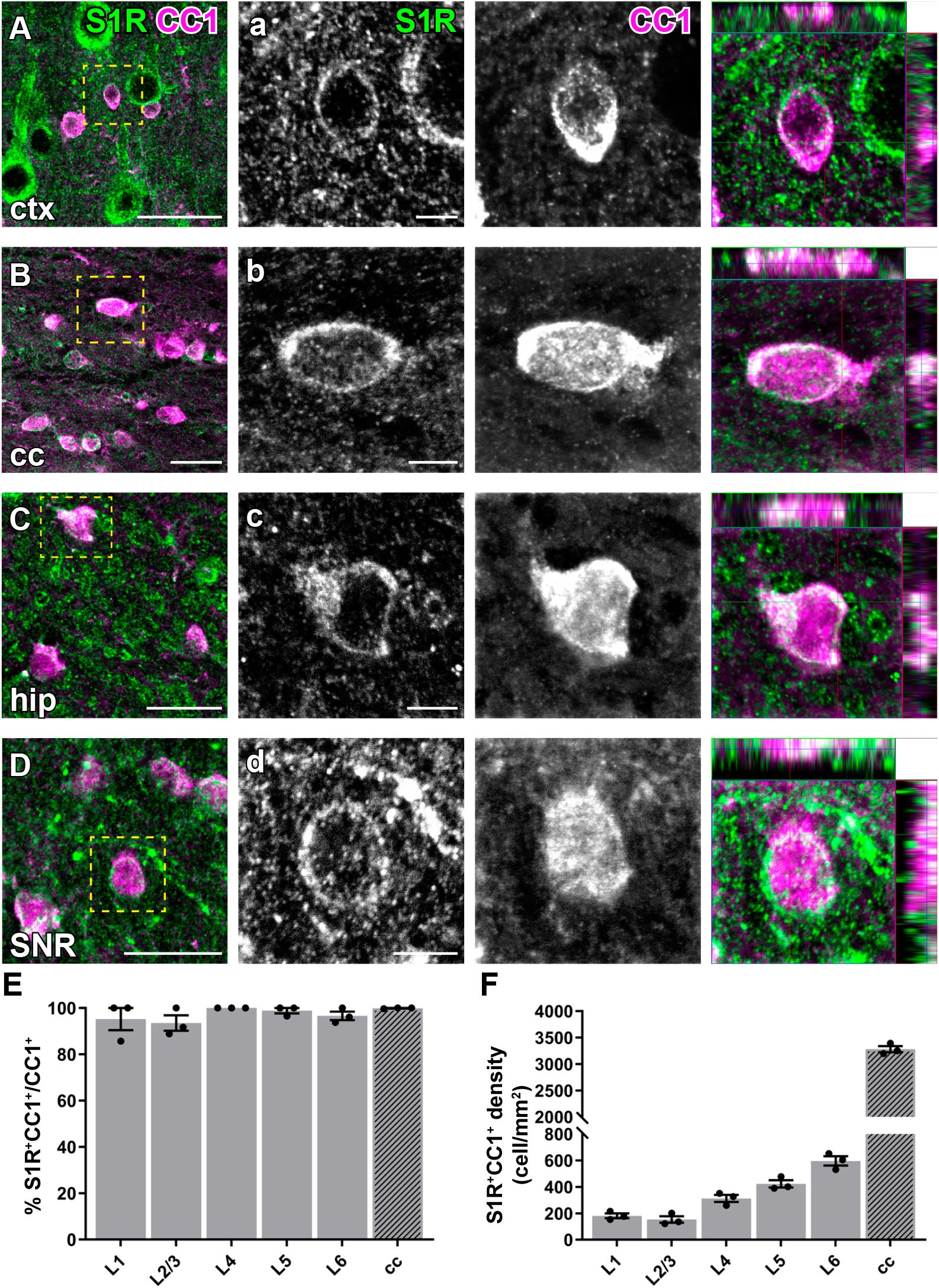
Immunohistochemical detection of S1Rs in forebrain oligodendrocytes. (**A-D**) S1Rs immunoreactivity in CC1^+^ oligodendrocyte in ctx (A), cc (B), hip (C), SNR (D). (**a-d**) Magnified images correspond to boxed areas in A-D. Arrowhead indicated S1R immunostaining in oligodendrocyte. The orthogonal views are shown in the rightmost images (**E-F**) Quantification of proportions (E) and densities (F) of S1R^+^CC1^+^ cells in different cortical layers (L1-L6) and cc. n = 3 mice. Scale bars = 20 µm in A-D, 5 µm in a-d.

To study the microglial expression of S1Rs, we combined Iba1 (a microglia marker) and S1R immunostaining. Like other glial cells, the majority of Iba1^+^ cells were immuno-labelled for S1R in all regions of the forebrain (Figure 7A-D). Quantification results even suggested that virtually all Iba1^+^ cells were co-expressing S1Rs, at least in the cortex and cc (Figure 7E-F).

**Figure 7.**
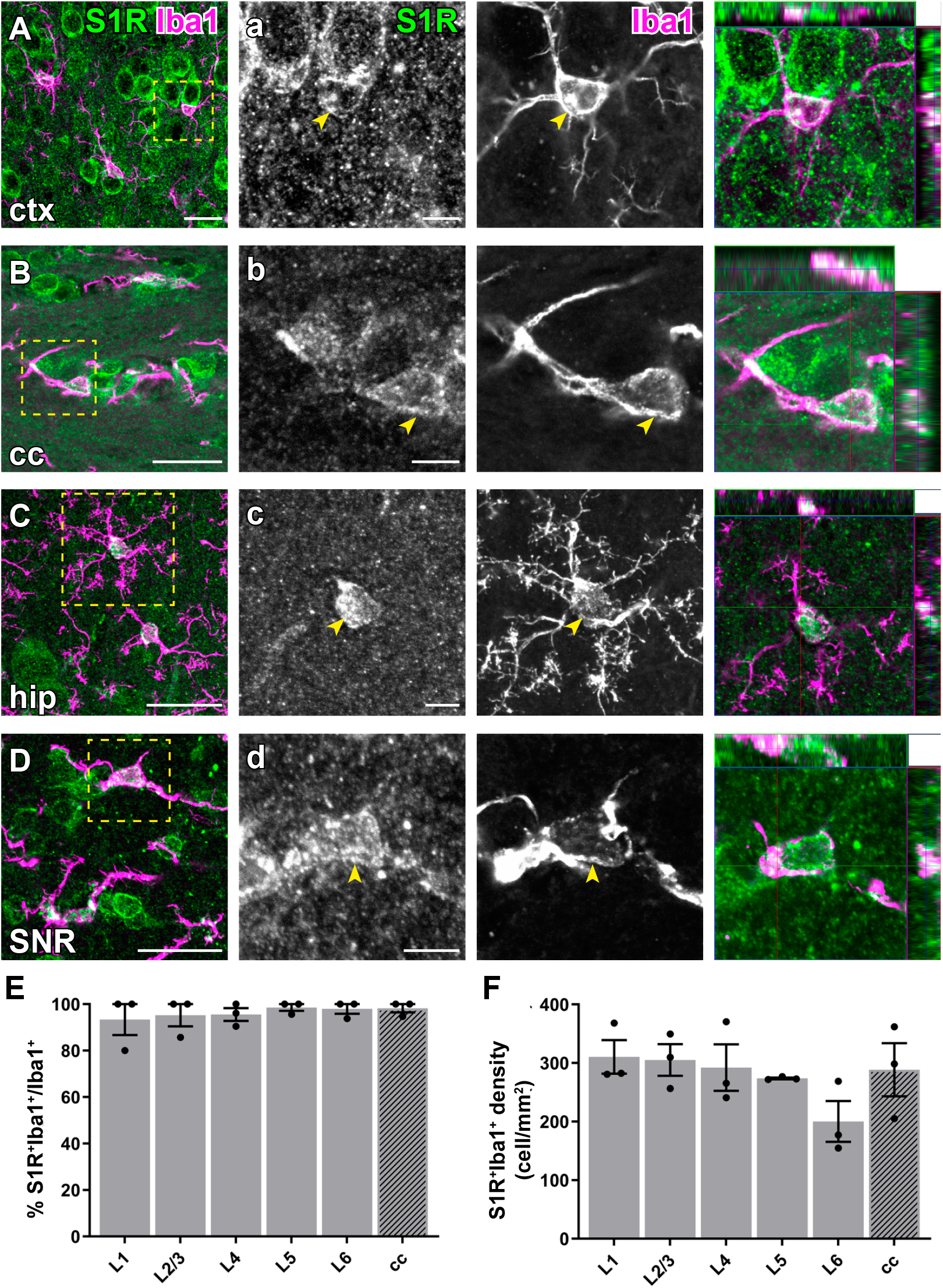
Immuno-labelling of S1Rs in microglia in the forebrain. (**A-D**) S1R immuno-labelling colocalized with Iba1 staining in ctx (A), cc (B), hip (C), SNR (D). (**a-d**) Magnified images corresponding to boxed regions in A-D. Arrowhead indicated overlapping of S1R and Iba1 immunostaining in microglia. The orthogonal images are presented at the rightmost. (**E-F**) Proportions (E) and densities (F) of S1R^+^Iba1^+^cells in different cortical layers (L1-L6) and cc. n = 3 mice. Scale bars = 20 µm in A-D, 5 µm in a-d.

Taken together, we were able to demonstrate that the majority of neurons and all types of glial cells express S1Rs based on the specific S1R immuno-labelling.

### S1Rs are widely expressed in the cerebellum and spinal cord

The AR^SDS^ protocol for S1Rs also generated specific immuno-labelling of S1Rs in the cerebellum (Figure 8A-B). We observed high expression in cell bodies and neurites of Purkinje neurons (confirmed by Calbindin staining) (Figure 8C). Further investigations with other cell type markers revealed that S1Rs were widely expressed in astrocytes including Bergmann glia (S100B^+^), microglia (Iba1^+^), OPCs (Pα^+^), and OLs (CC1^+^) in the cerebellum (Figure 8D-G).

**Figure 8.**
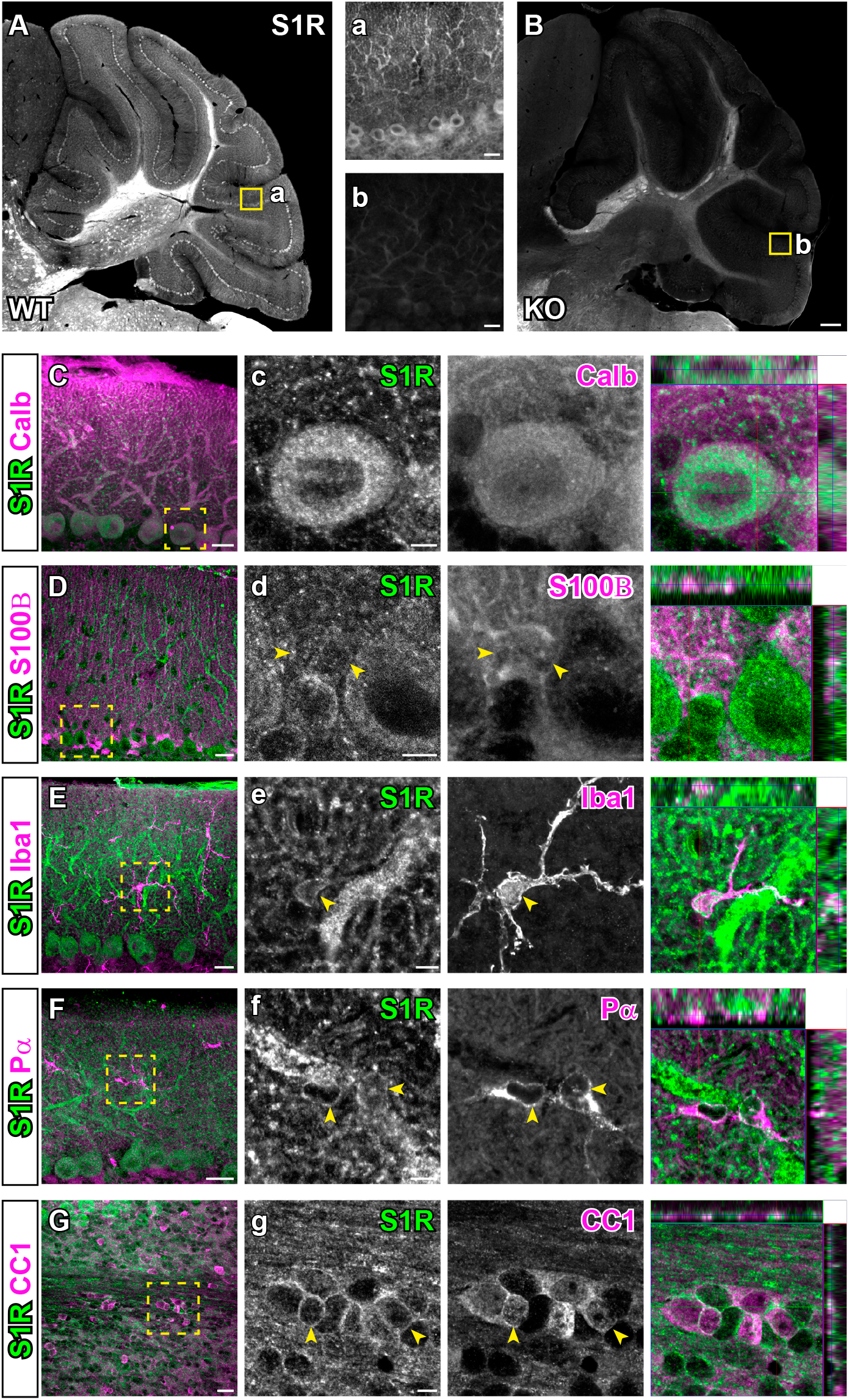
The expression of S1Rs in the cerebellum. (**A-B**) Sagittal sections of cerebellum were stained for S1R using the AR^SDS^ protocol. Specific immunostaining of S1Rs in WT cerebellum (A) is absent in S1R KO mouse (B). (**a-b**) Magnified views showing the molecular layer in the indicated regions of A and B. (**C-G**) Cerebellar slices were double-immunostained for S1Rs, calbindin^+^ (calb) (C), S100B^+^ (D), Iba1^+^ (E), Pα^+^(F) and CC1^+^(G), respectively. (**c-g**) The enlarged images of boxed area in C-G. Scale bars = 200 µm in A-B, 20 µm in a-b, C-G, and 5 µm in c-g.

Previous studies using Ab^Ruoho^ showed that high expression of S1R in motor neurons of the spinal ventral horn, however, the S1R expression in other cell types had not been mentioned. In the current work, we first tested the performance of the AR^SDS^ protocol for S1Rs in the spinal cord (Figure supplement 7A). We observed high expression of S1Rs in the ventral horn of WT mice, but with a strong background. Moreover, unspecific staining by the AR^SDS^ protocol could also be observed in the KO mice, preferentially in white matter myelin structures. Considering delipidation of myelin is widely used to decrease background for the detection of myelin proteins (Ishii, Fyffe-Maricich, Furusho, Miller, & Bansal, 2012; Jahn, Tenzer, & Werner, 2009), we tested a pre-treatment of spinal slices with 100% ethanol overnight (more than 16 h), followed by 1% SDS treatment. This modified protocol (AR^EtOH-SDS^, i.e. 100% ethanol + 1% SDS + Ab^#61994^; Figure 2A), largely reduced the background staining both in WT and KO mice (Figure supplement 7B). Therefore, we performed co-immunostaining for S1Rs and glial markers with the AR^EtOH-SDS^ protocol on spinal sections. We found that virtually all neurons in the spinal cord, though with even higher level in the ventral horn, were expressing S1Rs (Figure supplement 7C-D). Regarding the still relatively higher immunostaining background in the spinal cord compared to the brain, we quantified S1R immuno-positive cells in WT and KO spinal cords. We found that more than 90% of all types of glial cells either in the spinal grey matter or white matter of WT mice were immuno-labelled for S1R, whereas this proportion was no more than 5% in the KO mice (Figure supplement 8 and 9). Therefore, this AR^EtOH-SDS^ protocol still could be used to specifically detect S1R-expressing cells in the spinal cord. However, for yet unknown reasons, the myelin-like staining could still be seen in both groups of mice with the AR^EtOH-SDS^ protocol. To further investigate the unspecifically stained components, we combined myelin basic protein (MBP, a myelin marker) and S1R immunostaining. The unspecific S1R immuno-labelling did not fully overlap with MBP staining, but appeared to be in the inner layers of myelin sheaths (Figure supplement 7E-F). Thereby, this protocol is not suitable to study S1R expression in myelin in the spinal cord. More efforts are required to improve the AR of spinal slices for S1R immunostaining. Nevertheless, the current results from the AR^EtOH-SDS^ protocol demonstrated that the majority of neurons and glial cells in the spinal cord express S1Rs.

### Detection of S1Rs in the injured brain by immunohistochemistry

It has been suggested that S1R plays important role in the injured brain. However, the S1R expression pattern under neuropathology was not yet clear due to the lack of a reliable immunodetection method. Therefore, we evaluated the performance of the newly established S1R immuno-labelling AR^SDS^ protocol working on brains with acute brain injuries. We performed cortical stab wound injuries to adult mice which were analyzed at 3 days post injury (dpi). The IHC results showed that S1Rs in the (peri-) injured area were still well detected by the current protocol. We observed that the S1R expression level in the ipsilateral cortex did not show overt difference compared to the contralateral side, although the core injury area showed reduced S1R expression possibly due to loss of neurons (contra, Figure 9A, a_1_; ipsi, Figure 9A, a_2_). Moreover, S1Rs were also detected in activated microglia, astrocytes and OPCs in the injury-affected region (Figure 9B-D). Of note, we still did not find S1R immunostaining colocalized with GFAP in the main processes of astrocytes (Figure 9B, b). Compared to resting microglia in healthy brain, activated microglia showed stronger S1R expression, though its biological meaning remains to be elucidated (Figure 9C, c).

**Figure 9.**
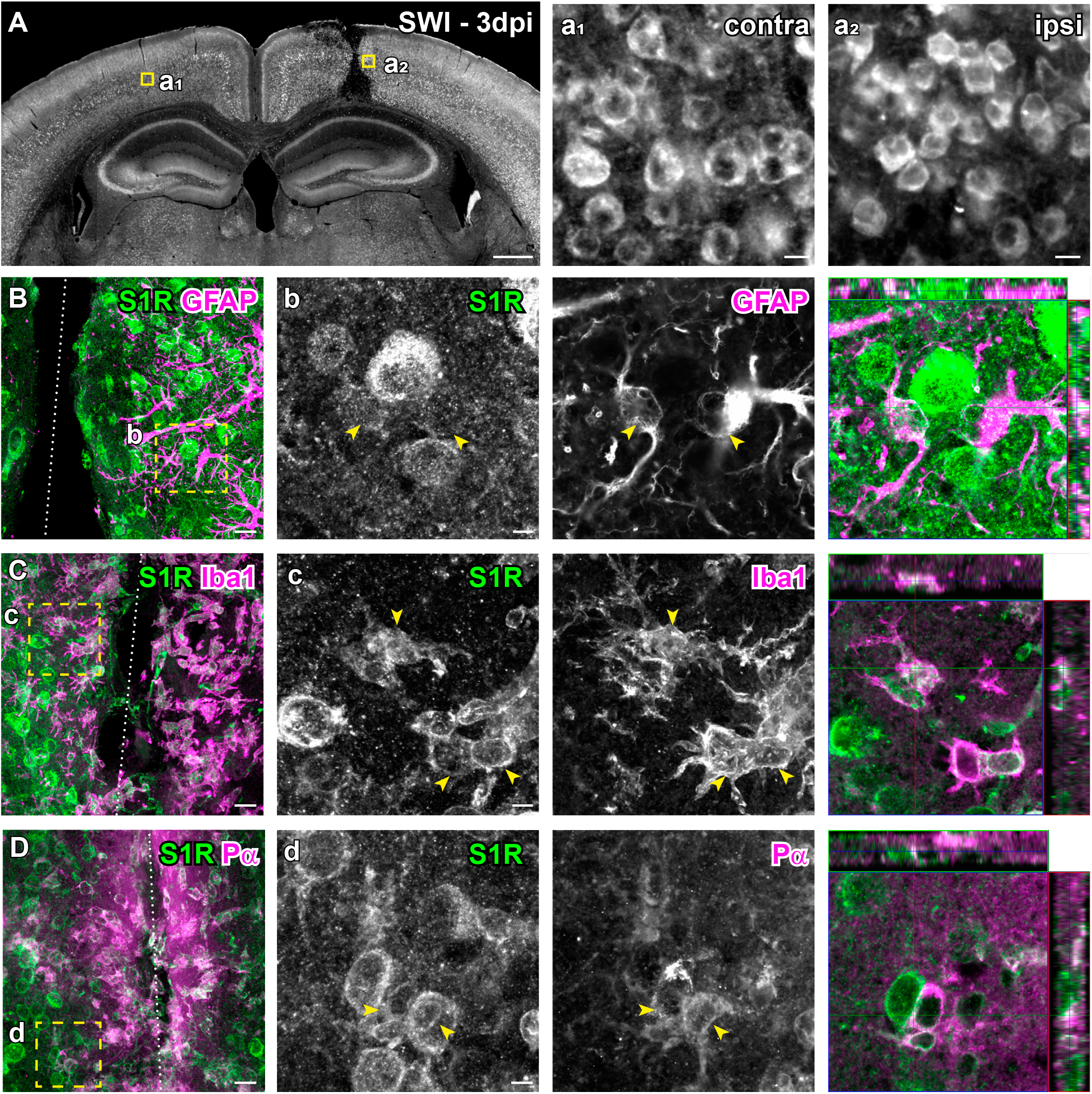
Immuno-labelling of S1Rs in the injured brain. (**A**) Overview of S1R immunostaining at 3 days post stab wound injury (3dpi). Ring-like structures of S1Rs in the contralateral (contra, a_1_) and ipsilateral (ipsi, a_2_) sides as indicated by the yellow boxes in A. (**B-D**) Injured brain sections were co-immunostained for S1Rs with various glial markers (GFAP, Iba1 and Pα). (**b-d**) The boxed areas in B-D are magnified. Activated astrocytes (GFAP^+^), microglia (Iba1^+^) and OPCs (Pα^+^) at the lesion site were expressing S1Rs enriched in the cell body. Scale bars = 500 µm in A, 10 µm in a_1_-a_2_, 20 µm in B-D, and 5 µm in b-d.

### Conditional deletion of S1Rs in the CNS *in vivo*

The newly established IHC protocol detecting S1Rs specifically enabled us to show that S1Rs are widely expressed in neurons and glial cells in the CNS. Therefore, the conditional knockout of S1Rs in cell types of interest *in vivo* would be a valuable tool to study S1R functions in the CNS. To achieve this goal, we generated a novel S1R flox mouse in which the exons 1-3 of *Sigmar1* are flanked by loxP sites (Figure supplement 10A). To test whether S1Rs could be specifically deleted in neurons *in vivo*, we crossed S1R flox mice to NEX-Cre knockin mice in which principal neurons express Cre to generate neuronal S1R cKO mice (NEX-Cre x S1R^fl/fl^) (Figure 10A). Control mice (NEX^wt/wt^ x S1R^fl/fl^) and cKO mice were analyzed at 10 w. We were able to show that S1R expression was drastically reduced in pyramidal neurons in the expected brain regions such as neocortex and hippocampus (Figure 10B-C). Quantification of S1R-expressing NeuN^+^ cells in the dorsal cortex confirmed that either in proportion (∼96% in ctrl, ∼5% in cKO) or in density (2900 cells/mm^2^ in ctrl, 156 cells/mm^2^ in cKO) S1R expression was largely ablated in NeuN^+^ cells in the neuronal S1R cKO mice (Figure 10D-E).

**Figure 10.**
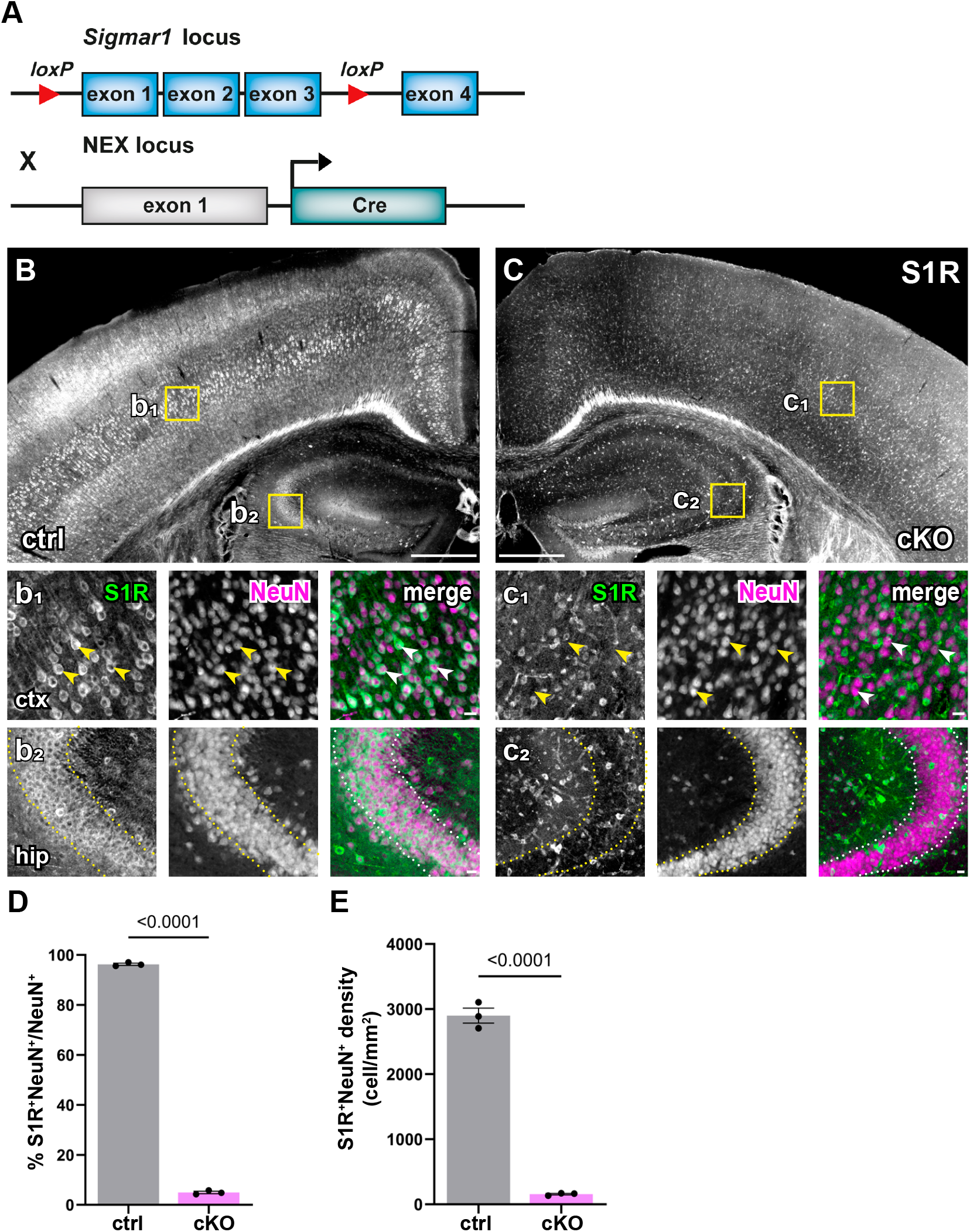
Successful knockout of S1Rs in principal neurons within the neocortex and hippocampus. (**A**) Scheme with transgenic structures of NEX-Cre x S1R^fl/fl^ mice used to delete S1Rs in principal neurons. (**B-C**) Overviews of S1R immunostaining in the dorsal brain of control (ctrl, B) and cKO (C) mice. (**b**_**1**_, **b**_**2**_, **c**_**1**_, **c**_**2**_) Magnified views (yellow boxes in B and C). Arrowheads indicate reduced S1R expression in cortical NeuN^+^ cells in cKO (c_1_) compared with ctrl (b_1_). Dotted lines indicate reduced expression of S1Rs in pyramidal neurons in the hippocampal CA2 region of cKO (c_2_) compared with ctrl (b_2_) mice. (**D-E**) Histograms highlighting decreased proportion and density of S1R^+^ cells in NeuN^+^ cells in cKO mice. n=3 mice per group. Scale bars = 500 µm in B-C, 50 µm in b_1_-b_2_, c_1_-c_2_.

To evaluate the temporally controlled deletion of S1Rs in targeted cell types of S1R flox mice, we crossbred S1R flox mice to <u>CX</u>3CR1-<u>C</u>reER<u>T</u>2 mice (CXCT^ct2/wt^ x S1R^fl/fl^) to generate microglia-specific S1R cKO mice (Figure 11A). These mice were injected with tamoxifen at 4 w and analyzed at 3 or 6 w post injection (wpi) (Figure 11B). Upon quantification of the immunostaining for Iba1 and S1R, we observed that in the dorsal cortex of cKO mice the proportion of Iba1^+^ microglia co-expressing S1R was reduced to 34.8 ± 2.9% at 3 wpi compared to ctrl (96.4 ± 3.2%). This level did not further decrease at 6 wpi (29 ± 6%) (Figure 11C-F). Thereby, we conclude that in adult mice the microglial S1R expression can be largely deleted within 3 weeks upon Cre induction.

**Figure 11.**
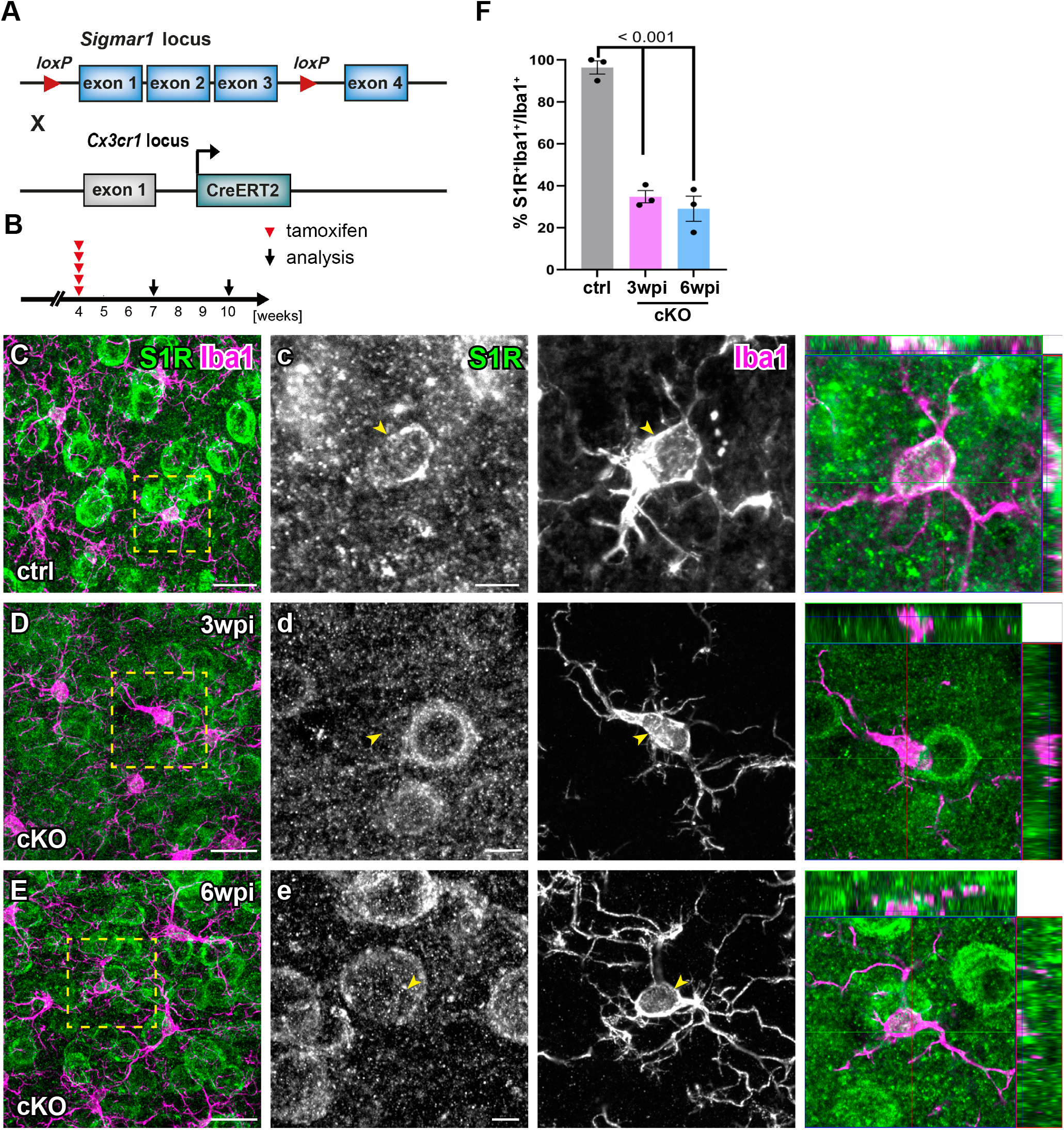
Successful deletion of S1Rs in microglia. (**A**) Schematic representation of double transgenic mice CX3CR1-CreERT2 x S1R^fl/fl^ used to conditionally delete S1Rs in microglia. (**B**) Experimental plan. All mice were injected with tamoxifen at 4 w. Immunostaining of S1Rs was performed at 3 w (3 wpi) or 6 w (6 wpi) post the first tamoxifen injection. (**C-E**) Double-staining of S1R and Iba1 in ctrl (C) and cKO (D-E) mice. (**c-e**) The boxed regions from B-D are magnified. The orthogonal views of S1R and Iba1 immuno-labelling are presented in the images at the right (**F**) Quantification S1R-expressing Iba1^+^ microglia in the ctx of ctrl and cKO mice. n=3 mice per group. Scale bars = 20 µm in C-E, 5 µm in c.

Taken together, the S1R flox mouse appears as a powerful tool to efficiently delete S1Rs in neurons and glial cells in the CNS *in vivo*.

## Discussion

Validation in KO animals is a golden standard to verify the specificity of antibodies for immunoblot and immuno-labelling (Laflamme et al., 2019). After screening of six commercial S1R antibodies using S1R KO mice, we identified Ab^#61994^ from Cell Signaling as a reliable antibody for specific detection of S1R using Western blotting of brain and spinal cord lysates. Ab^#61994^ is a newly produced rabbit monoclonal antibody, which has been used only in a few studies so far (six publications according to the Cell Signaling website). Recently, one study by Abdullah et al., verified the specificity of Ab^#61994^ to detect S1Rs in tissue lysate from mouse heart, but without showing the complete protein separation range of the immunoblot (Abdullah et al., 2020). Therefore, the current work provides further evidence that Ab^#61994^ is a reliable antibody specifically recognising S1Rs in immunoblot without generating bands at other molecular weight positions. However, four other S1R antibodies showed bands in protein samples from both WT and KO mice, indicating they are not working specifically for immunoblot. The antibody sc-137075 from Santa Cruz showed specific bands of S1Rs at the correct size, but also unspecifically bound to many proteins with different sizes other than S1Rs. Furthermore, we identified that only Ab^#61994^ could generate specific immunohistochemical staining of S1Rs with AR by 1% SDS. However, the other S1R antibodies failed to generate specific immunostaining *in vivo* under current tested conditions. Thus, our results suggest to cautiously re-evaluate previous studies using those antibodies for immunoblot, immunoprecipitation and/or immunohistochemistry of S1Rs.

The S1R has been discovered for over forty years. Intensive *in vitro* studies have revealed that S1Rs are serving as a pluripotent modulator of various cellular functions and are ligand-operated chaperons mainly localized on the ER membrane (Su et al., 2016). Recent transcriptomic studies of either bulk sequencing of purified CNS cells or single-cell sequencing all suggest that the S1R mRNA is widely detected in different cell types in the CNS (Consortium, 2020; Zhang et al., 2014). However, the spatial protein expression pattern of S1R is still difficult to conclude due to conflicting IHC results using antibodies generated by different research groups or commercial companies (Alonso et al., 2000; Hayashi & Su, 2004; Palacios et al., 2003). A recent study using S1R KO mice examined the specificity of the Ab^Rouho^ and several commercial S1R antibodies including Ab^sc-137075^ and Ab^42-3300^ for IHC in dorsal root ganglion (DRG), which suggested only Ab^Rouho^ could reliably label S1Rs in the DRG (Mavlyutov et al., 2016). Furthermore, Ab^Rouho^ was the only antibody validated by S1R KO mice for immuno-labelling of S1Rs in the CNS, though it did not work well for immunoblot (Mavlyutov et al., 2016; Mavlyutov et al., 2010). However, IHC studies using Ab^Rouho^ did not provide a clear S1R expression pattern at (sub)cellular levels in the CNS *in vivo*. In addition, Ab^Rouho^ is a custom-made antibody, hence, with limited availability to the research community.

In the current study the newly established IHC protocol (AR^SDS^ protocol) using Ab^#61994^ clearly revealed that S1Rs are mainly localized in the ER-like structure of CNS cells as suggested by *in vitro* studies. Notably, unlike the study using the Ab^Rouho^ (Mavlyutov et al., 2016) the current protocol demonstrated high expression levels of the S1R in the olfactory bulb, cerebral cortex, hippocampus and thalamus. Combining immunostainings with cell type-specific markers, we were able to show that in addition to neurons, as suggested by using Ab^Rouho^ (Mavlyutov et al., 2010), S1Rs are widely expressed in various glial cells in the CNS including astrocytes, OPCs, OLs, and microglia. Several AR methods such as heating with citrate buffer, microwave treatment, etc., have been tested to improve the immunostaining quality for the S1R in cultured cells (Hayashi, Lewis, Hayashi, Betenbaugh, & Su, 2011). In the current study, 1% SDS was used for the AR of the formaldehyde-fixed CNS tissue, substantially improving the S1R immunostaining. However, no AR was performed in studies using Ab^Rouho^ (Mavlyutov et al., 2016; Nakamura et al., 2019), which may explain their relatively fainter staining of S1Rs in the CNS compared to the results of the current protocols. Even more importantly, Ab^#61994^ is a monoclonal antibody produced by immortalized hybridoma cells, thereby ensuring its sustainable availability to the research community.

Although Ab^#61994^ displayed a very good capacity to specifically detect S1Rs in immunoblot and IHC, some drawbacks of using this antibody have to be considered. First, regarding the punctate background signals in the S1R KO brain, the signal/noise ratio of the current IHC protocol was not satisfactory to identify potential S1R expressions in fine structures such as the plasma membrane *in vivo*. Second, Ab^#61994^ can unspecifically bind to myelin-like structures in the spinal cord, brain stem and cerebellar white matter, hence limiting its application in white matter studies in certain CNS regions. Third, Ab^#61994^ did not work for IHC in brain tissues without SDS treatment, indicating that Ab^#61994^ may only recognize denatured S1R proteins. Therefore, it would be difficult to immunoprecipitate naïve S1Rs using Ab^#61994^. Further optimization of AR protocols for Ab^#61994^ as well as newly designed S1R antibodies would help to solve such problems.

Activation or inactivation of S1Rs by exogenous ligands both showed therapeutic effects for numerous neurological and psychological disorders. However, endogenous ligands of the S1R still remain unclear, hindering the understanding of patho-as well as physiological roles of receptor *in vivo* (Sałaciak & Pytka, 2022; Schmidt & Kruse, 2019). Loss-of-function experiments would be an ideal strategy to investigate functions of S1R. However, unlike the pharmacological effects of the exogenous ligands, S1R KO mice showed mild phenotypes in aging-related memory loss, cognitive impairments, motor defects, etc. (Couly et al., 2022). One possible explanation to such discrepancy can be a certain genetic compensatory machinery established during embryonic development that rescues the loss of S1R functions in global KO mice (El-Brolosy & Stainier, 2017). For example, hepatocyte-specific SIRT1 cKO mice develop a fatty liver which was not seen in global SIRT1 KO mice (Wang, Li, & Deng, 2010). Moreover, neither pharmacological intervention nor constitutive deletion of S1R could exclusively study S1R functions in specified cell types. To address such questions, we generated a novel Cre-dependant S1R conditional KO mouse (S1R flox). For the proof of concept, we showed that S1Rs could be successfully deleted in neurons or microglia mediated by Cre or CreER/tamoxifen systems *in vivo*. Regarding the broad expression of S1R in various cell types in and outside of the CNS, this novel S1R flox mouse will be a powerful tool to study cell-type specific functions of the S1R *in vivo*.

## Acknowledgement

We thank Daniel Schauenburg and colleagues for excellent animal husbandry and Frank Rhode for experimental assistance. This work was supported by grants from the Deutsche Forschungsgemeinschaft (Sino-German joint project KI 503/14-1 to WH and FK, SPP 1757, SFB 894 to FK); the Fritz Thyssen Foundation (10.21.1.021MN) and the Medical faculty of the University of Saarland (HOMFORexzellent2016) to WH; the BMBF (EraNet-Neuron BrIE) and the European Commission (EC-H2020 FET ProAct Neurofibres) to FK; the National Natural Science Foundation of China (81761138048) to HY. QL was supported by a PhD stipend from the Chinese Scholarship Council.

## Author contributions

WH conceptualized the project and designed experiments. WH and FK supervised the study. QL performed immunohistochemistry, immunoblot, qPCR and confocal imaging. QG performed RiboTag immunoprecipitation and qPCR. L-PF performed MACS. AS performed AxioScan imaging. QL analyzed data and generated figures. HY provided materials. WH and QL wrote the manuscript with inputs from all other authors.

