## Supplementary material for "Specific detection and deletion of the Sigma-1 receptor in neurons and glial cells for functional characterization *in vivo*": Figure supplement

### Supplementary figures

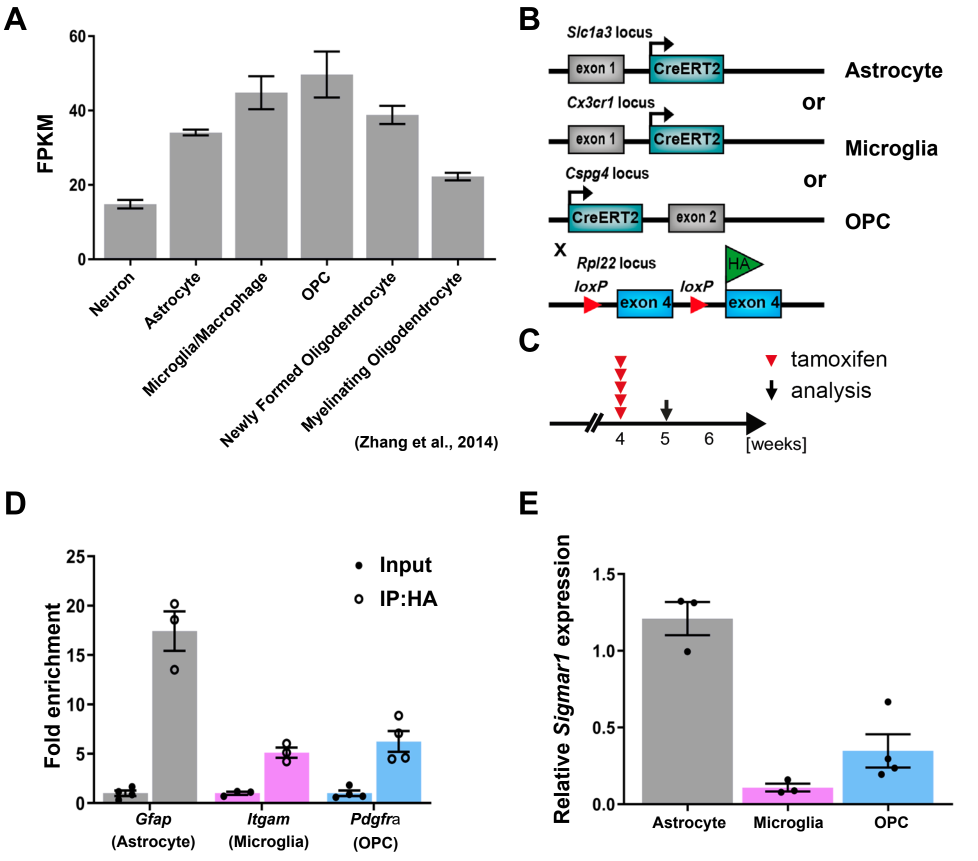


**Figure supplement 1. The mRNA expression of S1Rs in different glial cells in mouse brain.**

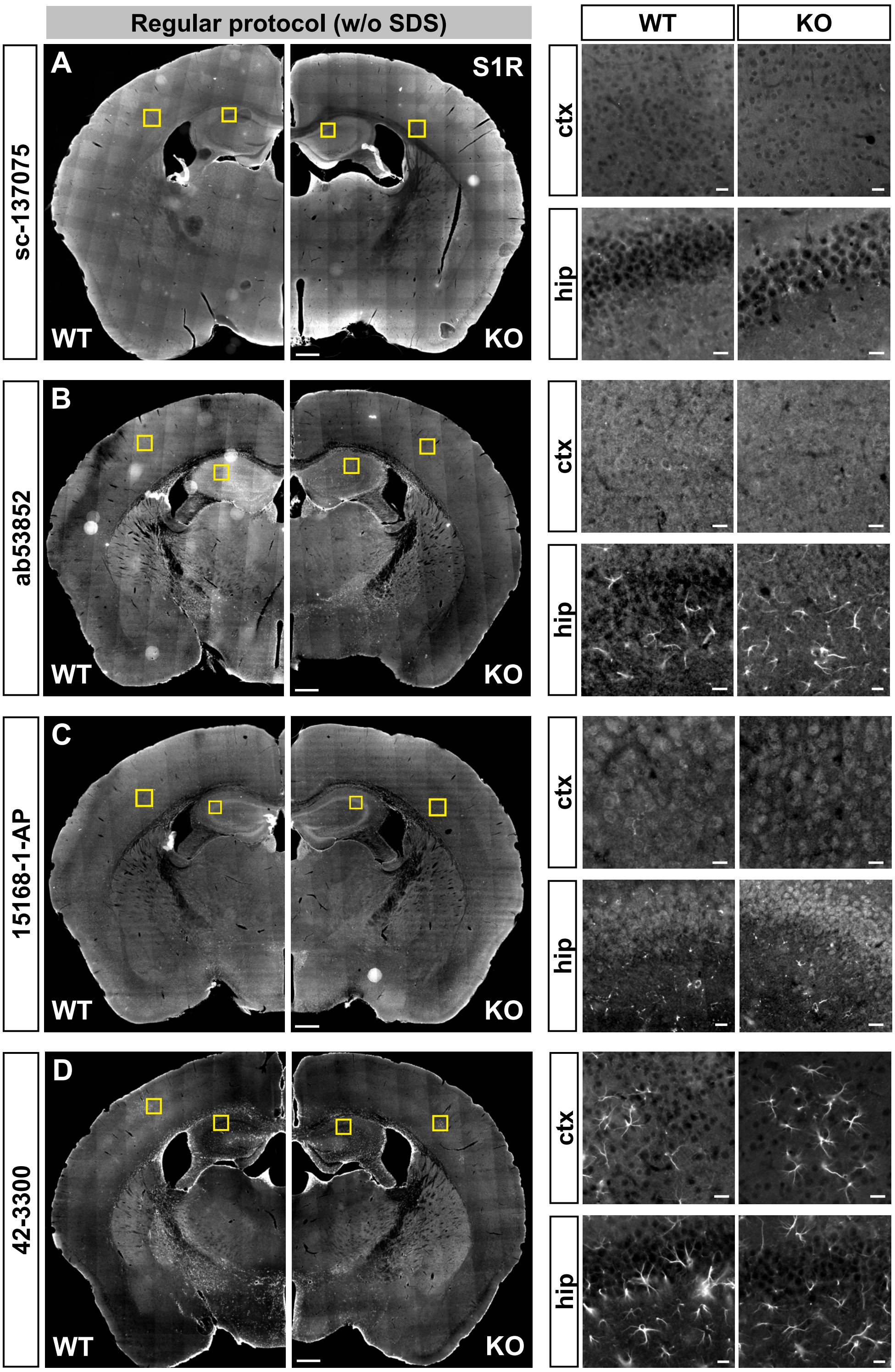


**Figure supplement 2. Immunostaining using the regular protocol with various commercial S1R antibodies.**

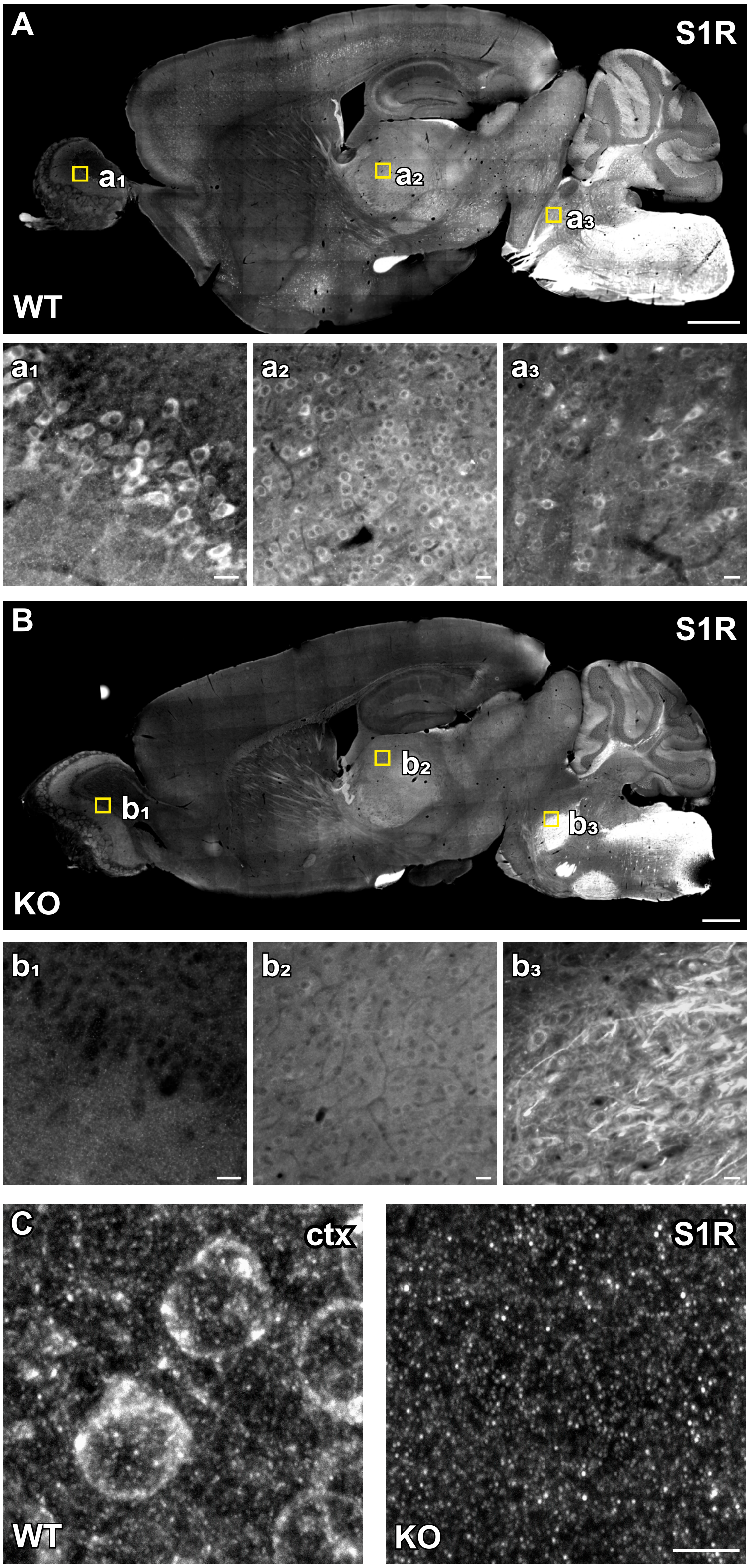


**Figure supplement 3. Overview of S1R immuno-labelling in the brain using the AR$^{SDS}$ protocol.**

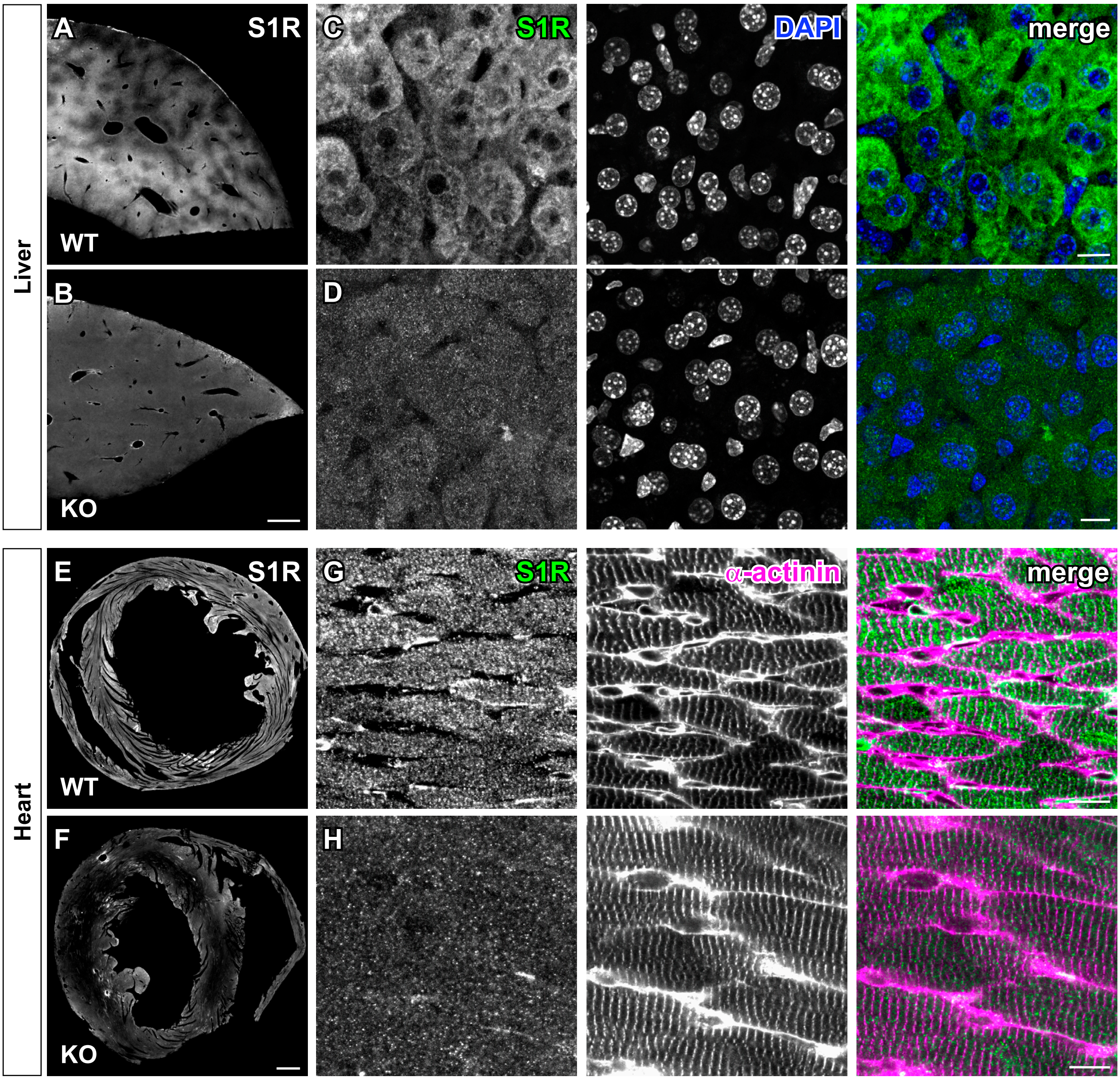


**Figure supplement 4. Specific detection of S1Rs in liver and heart.**

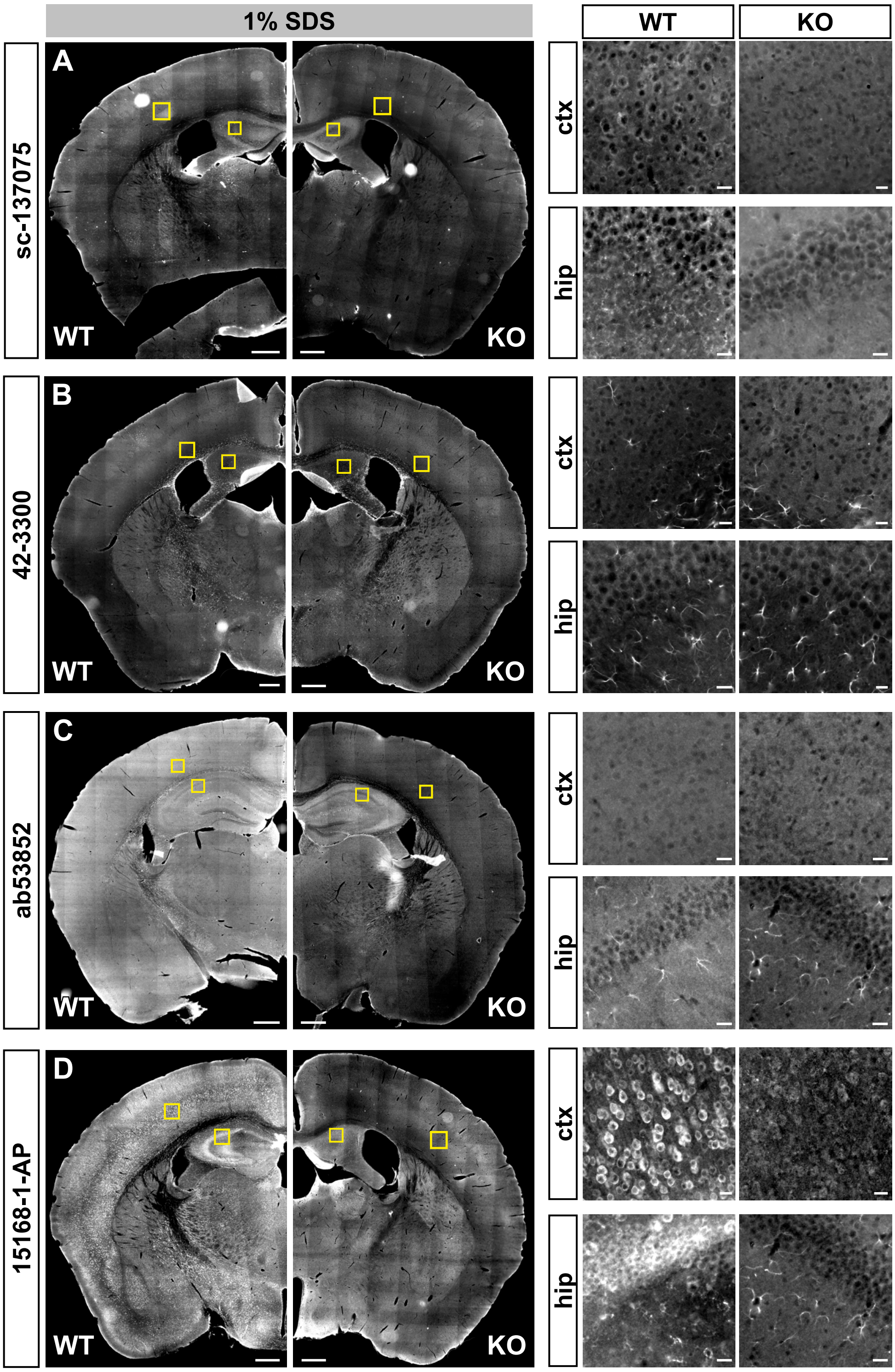


**Figure supplement 5. Performance of various commercial S1R antibodies in immunohistochemistry with SDS-antigen retrieval.**

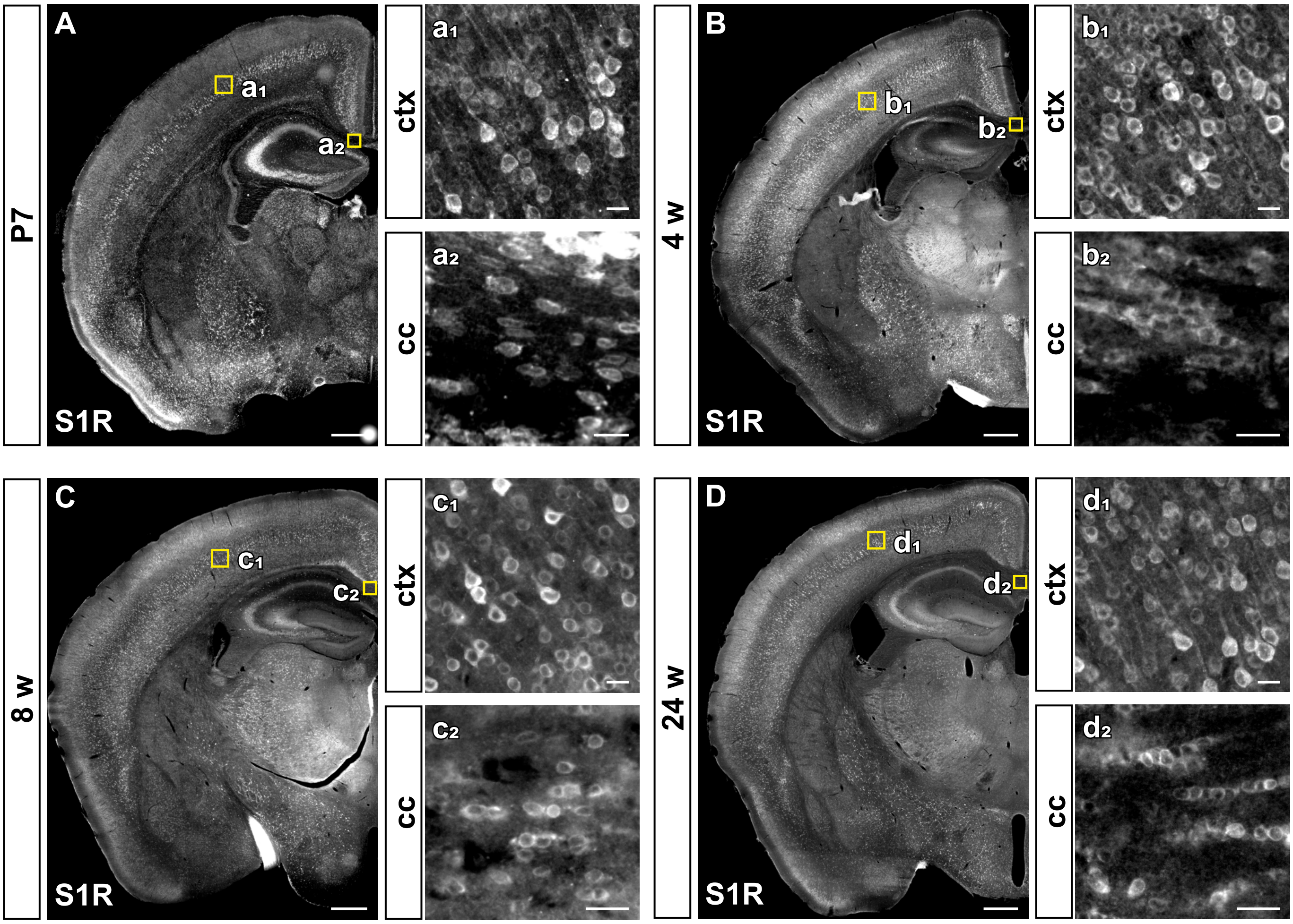


**Figure supplement 6. The expression pattern of S1Rs in the mouse brain at different ages.**

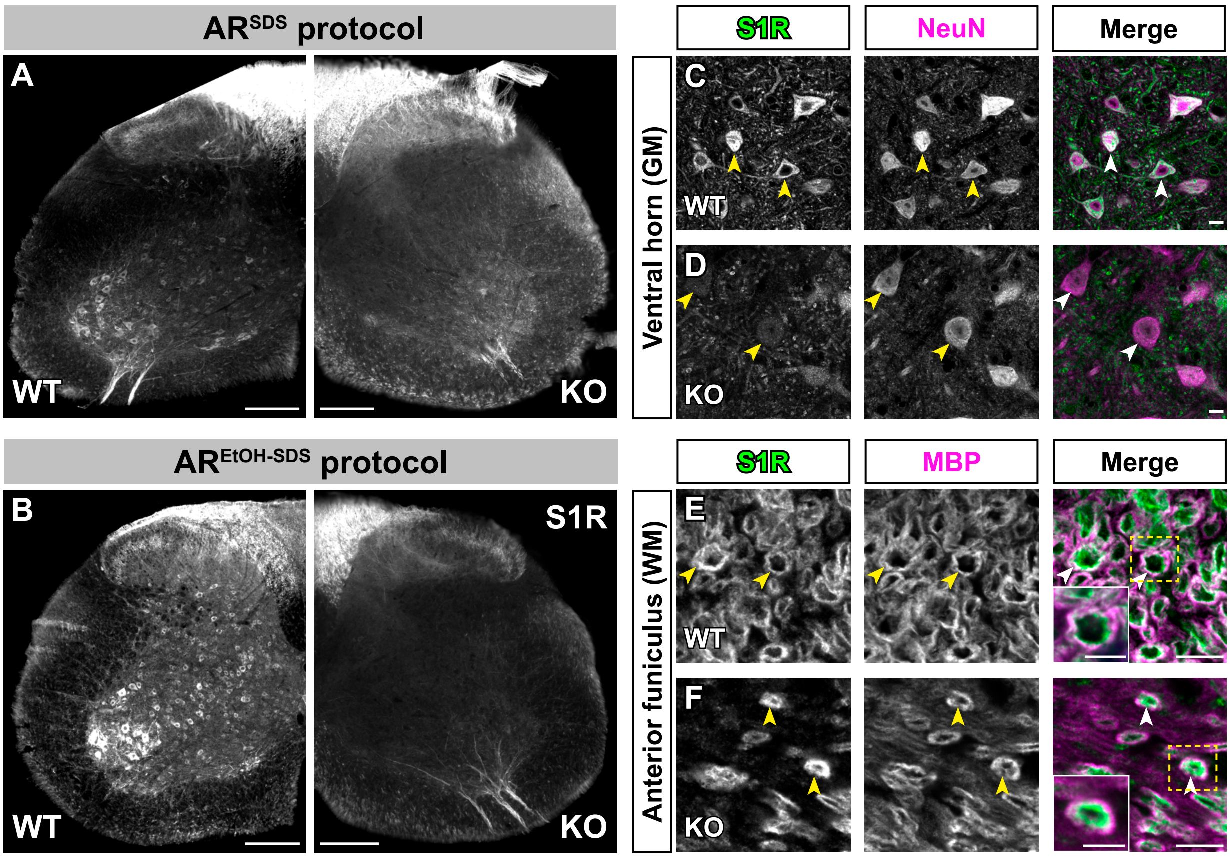


**Figure supplement 7. A modified AR$^{SDS}$ protocol to immuno-label S1Rs in mouse spinal cord.**

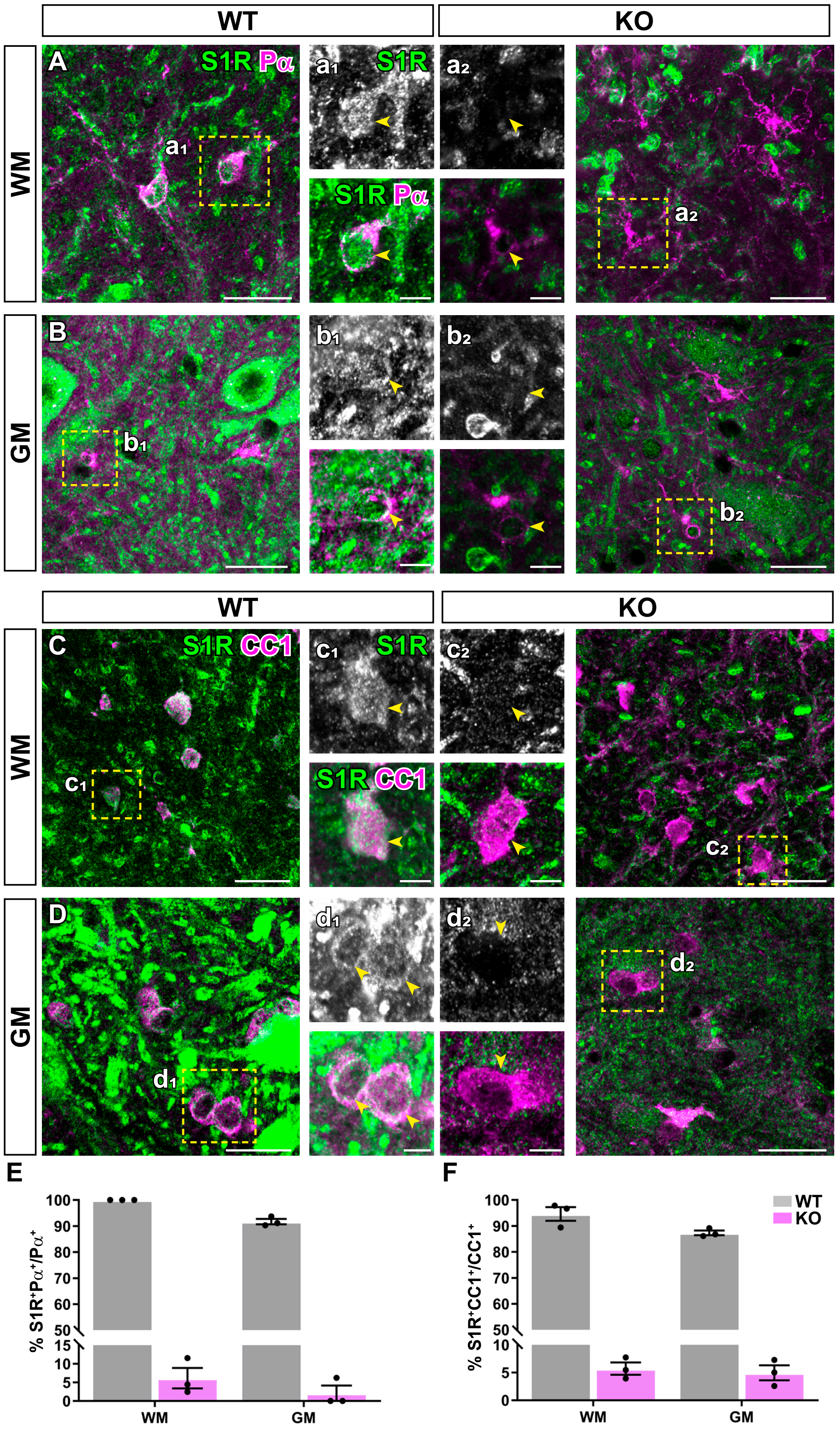


**Figure supplement 8. Specific detection of S1Rs in oligodendrocyte lineage cells in mouse spinal cord.**

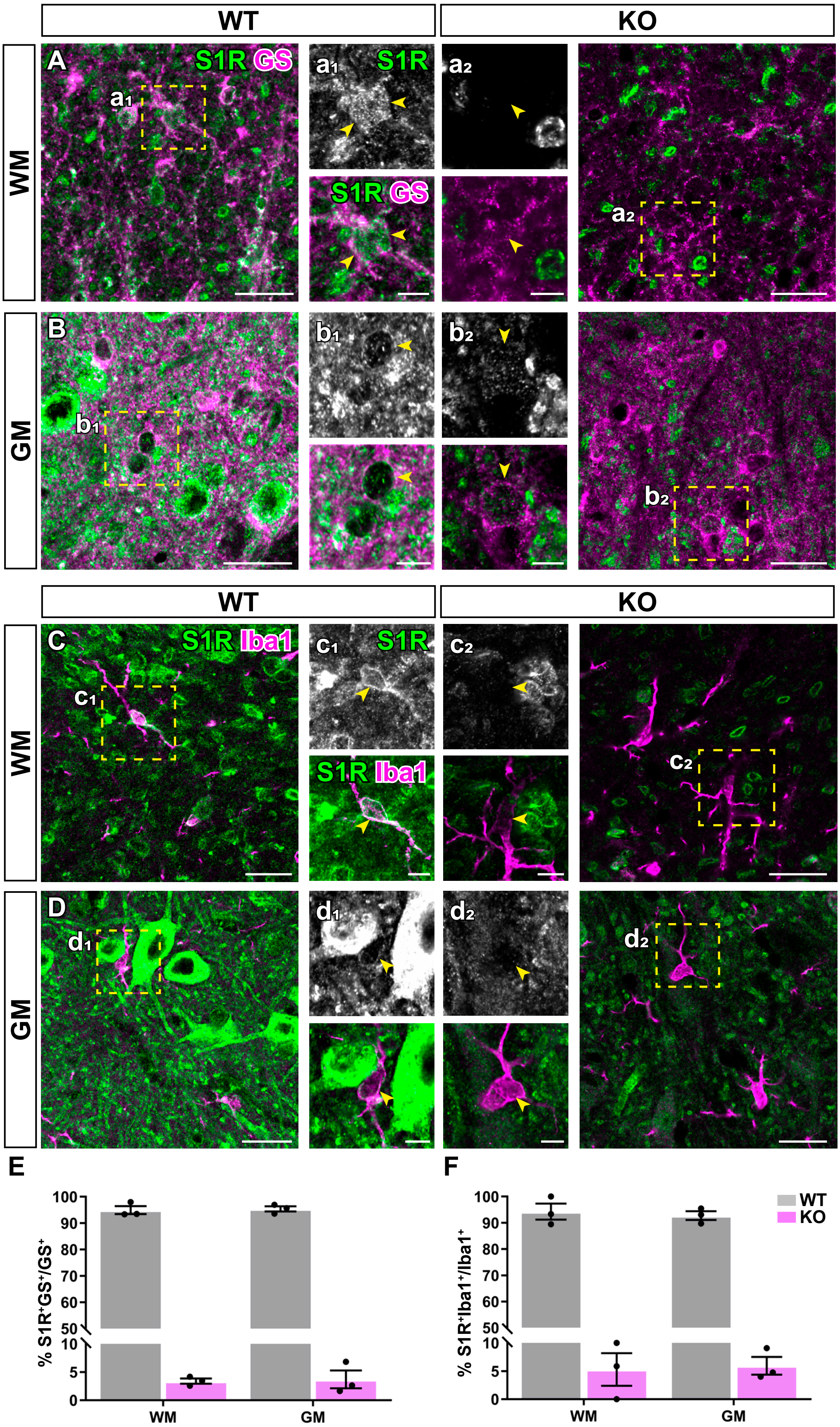


**Figure supplement 9. Specific immuno-labelling of S1Rs in spinal astrocytes and microglia.**

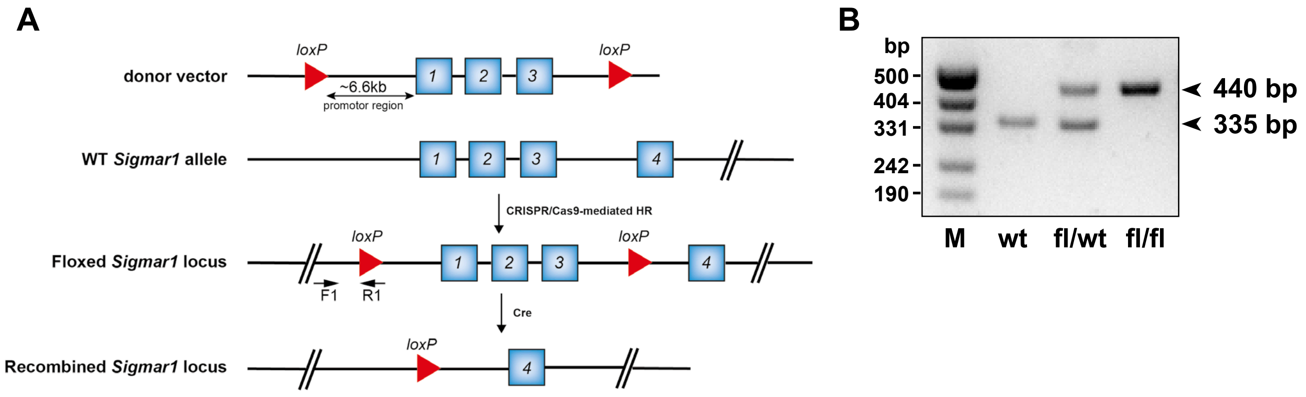


**Figure supplement 10. Generation of Cre-dependent *Sigmar1* flox mice.**

**Figure S1. The mRNA expression of S1Rs in different glial cells in mouse brain.** (**A**) Results from Brainrnaseq database (Zhang et al., 2014) show *Sigmar1* mRNA levels in different types of purified brain cells from pups at postnatal day 7 or 17 (only for OLs). FPKM: Fragments per kilo base of transcript per million mapped reads. (**B**) Transgenic schemes of three glial-specific CreERT2-expressing mouse lines crossbred with RiboTag mice to purify translated mRNA in astrocytes, microglia and OPCs, respectively. (**C**) Experimental plan. Mice were injected with tamoxifen at 4 weeks old and analyzed at 1 week after injection. (**D**) Enrichment of different glial cells were detected using specific marker genes expression level of input control and immunoprecipitated RNA by qPCR. (**E**) qPCR analysis comparing the relative translated *Sigmar1* mRNA expression level in astrocytes, microglia and OPCs, with normalization to input control. Each point represents one mouse.

**Figure S2. Immunostaining using the regular protocol with various commercial S1R antibodies**. (**A-D**) Overviews of brain sections from WT and S1R KO mice immunostained with S1R antibodies from Santa Cruz (sc-137075), Abcam (ab53852), Proteintech (15168-1-AP) and Invitrogen (42-3300), respectively. Representative images for boxed regions in the cortex and hippocampus (hip) are listed on the right side accordingly. Scale bars = 200 μm in A-D, 5 μm in magnified images.

**Figure S3. Overview of S1R immuno-labelling in the brain using the $AR^{SDS}$ protocol.** (**A-B**) Overviews of S1R immunostaining in sagittal mouse brain sections images showing a broad distribution of S1R from rostral to caudal in WT (B) mice which were largely devoid in S1R KO (C) mice. Some unspecific signals could be observed mainly in myelin tracts in the brain stem and spinal cord. ($b_1$-$b_3$, $c_1$-$c_3$) Magnified views showing representative images of olfactory bulb (**$b_1$, $c_1$**), thalamus (**$b_2$, $c_2$**) and medulla (**$b_3$, $c_3$**) from WT and KO mice. (**C**)The specific S1R immunostaining in WT mice. Punctate background signal of S1R immunostaining could be observed both in WT and KO mice. Scale bars = 1000 μm in A-B and 20 μm in $b_1$-$b_3$ and $c_1$-$c_3$, 5 μm in C.

**Figure S4. Specific detection of S1Rs in liver and heart.** With $AR^{SDS}$ protocol, we performed S1R staining on liver (**A-D**) and heart (**E-H**) from WT and S1R KO mice. α-actinin was used as the marker of cardiomyocytes. S1R immunostainings images with lower magnifications show clearly contract of WT and S1R KO mice. Scale bars = 500 μm in A-B, E-F, 10 μm in C-D, G-H.

**Figure S5. Performance of various commercial S1R antibodies in immunohistochemistry with SDS-antigen retrieval**. (**A-D**) Brain sections from WT and S1R KO mice were pre-treated with 1% SDS and immunostained by various S1R antibodies from Santa Cruz (sc-137075), Invitrogen (42-3300), Abcam (ab53852) and Proteintech (15168-1-

AP), respectively. Magnified images on the right side correspond to boxed regions from ctx and hip in A-D. Scale bars = 200 μm in A-D and 5 μm in magnified images.

**Figure S6. The expression pattern of S1Rs in the mouse brain at different ages.** Immunostaining of S1R showing similar S1R expression patterns in the brain at postnatal day 7 (P7, **A**), 4 week (w) (**B**), 8 w (**C**) and 24 w (**D**). Scale bars = 500 μm in A-D, 20 μm in $a_1$, $a_2$, $b_1$, $b_2$, $c_1$, $c_2$, $d_1$, $d_2$.

**Figure S7. A modified AR$^{SDS}$ protocol to immuno-label S1Rs in mouse spinal cord.** (**A-B**) Transverse spinal sections were immunostained with the specific S1R antibody (#61994). To improve staining quality, we modified the AR$^{SDS}$ protocol (A) with an additional delipidation step using 100% EtOH prior to 1% SDS treatment (B). (**C-D**) The double-immunostaining of S1R and NeuN in the ventral horn (grey matter, GM) of WT and S1R KO spinal cord. Some faint signals of S1R staining can be detected in NeuN$^+$ cells in the KO ventral horn. (**E-F**) The immunoreactivity of S1R and MBP (a myelin marker) in the anterior funiculus (white matter; WM) of WT (E) and S1R KO (F) mice. Magnified images represent boxed regions in E and F. Arrowheads indicate the localizations of S1R and MBP immuno-labelling. Scale bars = 200 μm in A-B, 10 μm in C-F, and 5 μm in magnified images.

**Figure S8. Specific detection of S1Rs in oligodendrocyte lineage cells in mouse spinal cord.** (**A-B**) The S1R immunoreactivity in Pα$^+$ OPCs both in spinal WM (A) and GM (B) of WT and S1R KO mice. (**$a_1$-$a_2$, $b_1$-$b_2$**) Magnified images correspond to areas in yellow boxes from A and B. Arrowheads indicate localizations of S1R and Pα immunostaining. (**C-D**) TheS1R and CC1 immunoreactivities both in spinal WM (C) and GM (D) of WT and S1R KO mice. (**c1-c2, d1-d2**) Magnified views in boxed areas from C and D. Arrowheads indicated localizations of S1R and CC1 immunostaining. (**E-F**) The quantification of the proportions of Pα$^+$ (E) or CC1$^+$ (F) cells in the spinal GM and WM of WT and S1R KO mice. Scale bars = 20 μm in A-D, 5 μm in $a_1$-$d_2$. n = 3 mice per group.

**Figure S9. Specific immuno-labelling of S1Rs in spinal astrocytes and microglia.** (**A-B**) The S1R immunoreactivity in GS$^+$ astrocyte both in spinal WM (A) and GM (B) of WT and S1R KO mice. (**a1-a2, b1-b2**) Magnified images correspond to boxed areas from A and B. Arrowhead indicated localization of S1R and GS immunostaining. (**C-D**) The S1R immunoreactivity in Iba1$^+$ microglia both in spinal WM (A) and GM (B) of WT and S1R KO mice. (**c1-c2, d1-d2**) Magnified views in boxed areas from C and D. Arrowheads indicated localizations of S1R and Iba1 immunostaining. (**E-F**) The proportions of GS$^+$ (E) and Iba1$^+$ (F) cells bearing S1R immunoreactivity in the spinal GM and WM of WT and S1R KO mice. Scale bars = 20 μm in A-D, 5 μm in $a_1$-$d_2$. n = 3 mice per group.

**Figure S10. Generation of Cre-dependent *Sigmar1* flox mice.** (**A**) Schematic drawing of Cas9/CRISPR-mediated generation of the *Sigmar1* flox mice. Arrows indicate the positions of primers for genotyping PCR. (**B**) Genotyping PCR result using primers F1 and R1 for wt, floxed *Sigmar1* heterozygous (fl/wt) and floxed *Sigmar1* homozygous (fl/fl) mice, respectively. The product sizes are indicated. M: DNA Ladders.
